# Characterization of the genetic composition and establishment of a core collection for the INERA Robusta coffee (*Coffea canephora*) field genebank from the Democratic Republic of the Congo

**DOI:** 10.1101/2023.06.15.544964

**Authors:** Lauren Verleysen, Robrecht Bollen, Jean-Léon Kambale, Ebele Tshimi, Benjamin Ntumba Katshela, Jonas Depecker, Valérie Poncet, Dieu-Merci Assumani, Filip Vandelook, Piet Stoffelen, Olivier Honnay, Tom Ruttink

**Author notes:** Correspondence: Corresponding Tom Ruttink.

## Abstract

Cultivation of Robusta coffee is likely to gain importance because of its high disease resistance and climate envelope, but substantial effort must be made to improve its cupping quality and other agronomic traits. Robusta coffee genetic resources conserved in field genebanks can play an important role in this context, but many of them are limited and poorly characterized. In this study, we aimed to characterize the genetic composition of the historically important but until recently neglected INERA Coffee Collection in Yangambi (Democratic Republic of the Congo) and to propose a Robusta core collection. We combined genome-wide GBS marker sets with a novel versatile multiplex amplicon sequencing (HiPlex) screening assay, for the genetic screening of the INERA Coffee Collection and created tools for future genetic screening. A set of 730 coffee shrubs in the collection were genotyped at 86 HiPlex loci and 263 unique genetic fingerprints were identified. Representative fingerprints were compared to ‘Lula’ cultivars, ‘Luki’ cultivars, Congolese subgroup A, and wild genotypes growing in the rainforest of the Yangambi region. The majority of the unique genetic fingerprints were identified as ‘Lula’ cultivars but also four Congolese subgroup A and nine wild genotypes were found in the collection. Twenty-five percent of the unique genetic fingerprints had a hybrid identity which could result from seed-based propagation of the collection or natural cross-pollination. Moreover, eight unique genetic fingerprints were frequently identified as a parent and these may correspond to the ancient selected mother plants, or direct descendants thereof, of which seed material was widely distributed from 1930 to 1960 for the creation of coffee plantations. Finally, to delineate core collections we tested the maximization strategy and genetic distance method for the INERA Coffee Collection, and proposed i) a core collection conserving the loci diversity of the collection, *i.e.* 100 accessions and ii) a core collection holding the highest genetic distances between the accessions, *i.e.* 10 accessions. Our study provides a method for the genetic characterization of Robusta coffee collections in general and contributes to the revaluation and exploitation of the Robusta coffee genetic resources in the Democratic Republic of the Congo in particular.

## INTRODUCTION

Coffee is the world’s most widely consumed hot beverage and the second-most exported product from developing countries (Pendergast, 2009; Lashermes, 2018). Coffee belongs to the *Rubiaceae* family, and to the genus *Coffea* that comprises 131 species (Davis & Rakotonasolo, 2021; Stoffelen et al., 2021) of which only *Coffea arabica* L. (Arabica coffee) and *C. canephora* Pierre ex A. Froehner (Robusta coffee) are cultivated at a commercial scale. *C. canephora* has the widest native distribution range among all *Coffea* species, ranging from West Africa through Cameroon, Central African Republic, Republic of the Congo, the Democratic Republic of the Congo (DRC), Uganda, northern Tanzania down to northern Angola (Cubry et al., 2013). Within its native distribution range, two major origin groups were previously identified, namely the Congolese and Guinean groups (Montagnon et al., 1992; Dussert et al., 1999; Cubry et al., 2013). The Congolese group was further subdivided into seven subgroups: group A in Gabon, the Republic of the Congo and western DRC, group E in the DRC, group C in Cameroon and the western Central Africa region, group B in eastern Central African Republic, group O in Uganda (Gomez et al., 2009), and two recently described groups G in Angola, and R in southern DRC (Merot-L’anthoene et al., 2019). The Guinean group currently corresponds to group D. Of these eight origin groups, materials derived from mainly Congolese subgroup E and subgroup A or the Guinean group D are supposed to be used for *C. canephora* cultivation (Leroy et al., 1993; Montagnon et al., 1998a; Oliveira et al., 2018).

*C. canephora* was initially cultivated at a small scale in the late 19^th^ century in Gabon, Angola, Uganda, and the Sankuru region of the DRC (Durand et al., 1898; Chevalier, 1929; Montagnon et al., 1998a; Vanden Abeele et al., 2021). At that time, Arabica coffee cultivation in Asia was threatened by leaf rust disease, and plant hunters were searching for alternative coffee species from Africa. After several unsuccessful attempts to introduce novel species like *Coffea liberica*, Linden launched in 1900 a robust, pest-resistant, and productive coffee species under the name ‘*Coffea robusta*’. These genetic resources were introduced in the trials of the Java Coffee Research Station, which became the first important breeding and distribution center of Robusta (Ferrão et al., 2019). From 1910 on, Robusta was promoted and distributed as an important colonial cash crop, further stimulating Robusta coffee research and breeding activities and leading to the establishment of the Lula Coffee Research Station in the DRC, which was later integrated in the *Institut National pour l’Etude Agronomique du Congo Belge* (INEAC, after the independence of the DRC, the institute was renamed to INERA). In 1927, the Yangambi Research Station was established less than 100 km from the INERA Coffee Research Station in Lula, starting with a coffee research program focused on Robusta material derived from the Java Coffee Research Station and the INERA Coffee Research Station in Lula. Subsequently, the INERA Coffee Collection in Yangambi was enriched with other wild and cultivated material from the DRC (*e.g.* from the INERA Coffee Collection in Luki), and abroad, bringing *C. canephora* genetic resources from different origin groups together. From 1930 until 1960, the Yangambi Research Station (meanwhile also part of the INERA), was taking the lead in Robusta breeding in the DRC, and was distributing ‘Lula’ and ‘INEAC’ elite breeding lines worldwide (Coste et al., 1955; Montagnon et al., 1998b). In 1951-1952, seven mother plants with improved pest resistance, productivity, and quality, were created and their seeds were mixed to form a standard seed material blend that was widely distributed for the creation of plantations (Capot, 1962). With its large number of *C. canephora* genetic lines (wild accessions and cultivars) and its coffee research program, the Yangambi Research Station became the most important *C. canephora* selection center by 1950 (Montagnon et al., 1998b).

The once very rich INERA Coffee Collection in Yangambi did not escape the many difficulties the DRC has faced during the last decades and was decimated due to lack of appropriate care and funding (Stoffelen et al., 2019). Since 2016, initiatives have been undertaken to rehabilitate the INERA Coffee Collection in Yangambi and part of those efforts concern (genetic) characterization of the plant material. The collection is, especially since 2020, further enriched with numerous new accessions with a wild and cultivated origin collected from several regions within the DRC. It is currently not known how many and which of the ‘Lula’ and ‘INEAC’ elite breeding lines and other wild and local cultivated material from the original INERA Coffee Collection in Yangambi remain and whether they still correspond to the plant material currently grown in the field genebank. A preliminary survey of the collection management revealed several issues. First, inconsistencies were found between the field maps of the field genebank and the labeled accessions present on the field, and many accessions were missing their original plant label. Second, a broad phenotypic diversity for various morphological and agronomical traits were observed within plots that were assumed to contain clonally propagated, *i.e.*, genetically identical plant material, suggesting incorrect labeling (based on field observations 2020-2021, data not shown). Last, years of multiplication of accessions through sexual propagation (*i.e*. through seedlings rather than cuttings) and open pollination resulted in the hybridization of the initial accessions rather than maintaining unique lines. Vanden Abeele et al. (2021) provided a first exploratory screening of 45 coffee shrubs from the INERA Coffee Collection in Yangambi using Simple Sequence Repeat (SSR) markers and found that the majority of those accessions, which are referred to as Lula varieties, are presumably originated from the Coffee Research Station in Lula. In addition to these Lula varieties, that study identified several rare cultivars originating from the North Kivu, Orientale province, and Equateur province, and four Petit Kwilu cultivars, presumed to originate from western DRC, Republic of the Congo, and Gabon. Vanden Abeele et al. (2021) could also identify two wild genotypes originating from the Ituri and Tshopo provinces (DRC).

Establishing a core collection is key to the future sustainable and effective conservation management and use of the present INERA Coffee Collection in Yangambi, which highly likely contains valuable cultivated and wild genetic resources for coffee production and breeding. A core collection is a subset of the entire collection of germplasm (seeds, plants, or tissues) of a particular species that captures the most diversity with minimal redundancy (Brown, 1989). Currently, there are two complementary, commonly used strategies to construct a core collection: i) the maximization (M) strategy, which focuses on selecting the most diverse loci to maximize the genetic diversity of the core collection and ii) the genetic distance method, which aims to select the most diverse plant material within a collection to maximize the genetic distance between the entries of the core collection (Gu et al., 2023). Leroy et al. (2014) used these two core collection strategies to propose core collections of the genetic resources of *C. canephora* based on 565 *C. canephora* accessions collected from the Ivory Coast, Uganda, the DRC, and French Guyana and characterized them with 13 SSR markers. Using three different core sizes (12, 24, and 48 entries), they proposed seven core collections that can be used as a valuable tool for diversity management and preserving the genetic resources of *C. canephora*, or it can serve as a solid basis for breeding programs. By combining the M strategy and genetic distance method, Leroy et al. (2014) created an optimal core collection containing 77 accessions, which can be effectively utilized in research centers and in the context of improving coffee production through breeding efforts.

In this study, we genetically characterized 730 coffee shrubs carefully selected from the pre-2020 INERA Coffee Collection in Yangambi (hereafter referred as ‘Yangambi coffee collection’) with the aim to (i) develop a screening assay based on multiplex amplicon sequencing (HiPlex) to identify unique genetic fingerprints; (ii) genetically characterize this collection and classify the 730 individuals in unique genetic groups; (iii) specifically explore the presence of ‘Lula’ cultivars, cultivars from the INERA Research Station in Luki, Congolese subgroup A, and local wild coffee genotypes in the collection based on genetic similarity to reference samples; and (iv) propose representative core collections. The molecular markers were developed in several subsequent steps. First, a set of diverse genotypes (n=218, ‘Discovery Panel’) was compiled based on the field maps and plant labels and genotyped using genome-wide molecular markers (Genotyping-by-Sequencing) to identify unique genetic fingerprints. Second, a high-throughput screening assay based on multiplex amplicon sequencing (HiPlex) was created with sufficient genetic resolution to discriminate the unique genetic fingerprints. After validation, the HiPlex assay was used to comprehensively screen the Yangambi coffee collection (n=730, ‘Screening Panel’) and to identify all unique genetic fingerprints. Then, population structure and kinship were analyzed in three steps. First, the unique genetic fingerprints of the Yangambi coffee collection were compared to reference material of the Congolese subgroup A, previously described by Merot-L’anthoene et al. (2019) and wild genotypes growing in the local rainforest of the Yangambi region, previously described by Depecker et al. (2023) and assigned to Congolese subgroup BE by Merot-L’anthoene et al. (2019), to estimate the relative abundance of plant material of these origins in the collection. Second, fastSTRUCTURE and principal component analysis (PCA) were used to reveal population structure in the Yangambi coffee collection. Third, a parentage analysis was used to investigate what part of the population structure results from crosses and to delineate which genotypes were preferentially used for seed-based propagation. Finally, the M strategy and genetic distance method were tested for the Yangambi coffee collection to propose complementary core collections.

## MATERIAL & METHOD

### Plant material used for genotyping

Leaf material from 730 coffee shrubs was collected from the INERA Coffee Collection in Yangambi, hereafter referred to as the ‘Yangambi coffee collection’ (see details in Supplemental Table S1). Genomic DNA was extracted from 20-30 mg dried leaf material using an optimized cetyltrimethylammonium bromide (CTAB) protocol adapted from Doyle & Doyle (1987). DNA quantities were measured with the Quantifluor dsDNA system on a Quantus Fluorometer (Promega, Madison, USA). Of these 730 samples, 218 were subjected to GBS and all 730 samples were subjected to the HiPlex assay (see below). Genomic DNA extracts of 235 wild coffee shrubs collected from the local rainforest in the Yangambi region were obtained from Depecker et al. (2023). Genomic DNA extracts of 14 herbarium coffee samples collected from the INERA Coffee Collection in Luki were supplied by Meise Botanic Garden, Belgium, hereafter referred to as ‘Luki’ cultivars. All samples collected from the local rainforest and from the INERA Coffee Collection in Luki were subjected to the HiPlex assay. Whole genome shotgun (WGS) sequencing data of a wild sample from Republic of the Congo (accession 20708) and a cultivated sample from Togo (accession 20723), previously described by Merot-L’anthoene et al. (2019), were retrieved from Sequence Read Archive (SRA) (Tournebize et al., 2022; PRJNA803612), and were used as reference for the Congolese subgroup A.

### Genotyping-by-Sequencing and read data processing

Using GBS, we first screened a panel of 218 samples (hereafter referred to as ‘Discovery Panel’), representing the assumed genetic diversity based on documented background knowledge of the Yangambi coffee collection (field maps and plant labels), here referred to as ‘documented identities’. Then, following Depecker et al. (2023), GBS libraries were prepared using a double-enzyme GBS protocol adapted from Elshire (2011) and Poland et al. (2012). In short, 100 ng of genomic DNA was digested with PstI and MseI restriction enzymes (New England Biolabs (NEB), Ipswich, USA), and barcoded and common adapters were ligated with T4 ligase (NEB) in a final volume of 35 µL. Ligation products were purified with 1.6x MagNA magnetic beads (GE Healthcare Europe, Machelen, BE) and eluted in 30 µL TE. Of the purified DNA eluate, 3 µL was used for amplification with Taq 2x Master Mix (NEB) using an 18 cycles PCR protocol. PCR products were bead-purified with 1.6x MagNA, and their DNA concentrations were quantified using a Quantus Fluorometer. The library quality and fragment size distributions were assessed using a QIAxcel system (Qiagen, Venlo, NL). Finally, equimolar amounts of the GBS libraries were pooled, bead-purified, and 150 bp paired-end sequenced on an Illumina HiSeq-X instrument by Admera Health (South Plainfield, USA).

Reads were processed with a customized script available on Gitlab (https://gitlab.com/ilvo/GBprocesS). First, the quality of sequence data was validated with FastQC v0.11 (Andrews, 2010) and reads were demultiplexed using Cutadapt v2.10 (Martin, 2011), allowing zero mismatches in barcodes or barcode-restriction site remnant combination. Next, the 3’ restriction site remnant and the common adapter sequence of forward reads and the 3’ restriction site remnant, the barcode, and the barcode adapter sequence of reverse reads were removed based on sequence-specific pattern recognition and positional trimming using Cutadapt. After trimming the 5’ restriction site remnant of forward and reverse reads using positional trimming in Cutadapt, forward and reverse reads with a minimum read length of 60 bp and a minimum overlap of 10 bp were merged using PEAR v0.9.11 (Zhang et al., 2014). Merged reads with a mean base quality below 25 or with more than 5% of the nucleotides uncalled and reads containing internal restriction sites were discarded using GBprocesS. Finally, merged reads were aligned to the *C. canephora* reference genome sequence (Denoeud et al., 2014) with the BWA-mem algorithm in BWA v0.7.17 (Li, 2013) with default parameters. Alignments were sorted, indexed, and filtered on mapping quality above 20 with SAMtools 1.10 (Li et al., 2009). Next, high-quality GBS loci and Stack Mapping Anchor Points (SMAPs) were identified in the mapped reads using the SMAP *delineate* module within the SMAP package v4.4.0 (Schaumont et al., 2022) with *mapping_orientation* ignore, *min_stack_depth* 4, *max_stack_depth* 400, *min_cluster_depth* 8, *max_cluster_depth* 400, *completeness* 90, and *min_mapping_quality* 20.

### SNP calling

Single nucleotide polymorphisms (SNPs) were called with GATK (Genome Analysis Toolkit) Unified Genotyper v3.7.0. (McKenna et al., 2010). SNP calling was the same for GBS and HiPlex (see below). Only SNPs within high-quality GBS loci as identified with the SMAP *delineate* module, for GBS read data, or within the 86 high-quality HiPlex loci, for HiPlex read data, were retained. SNPs were filtered using the following parameters: *min-meanDP* 30, *mac* 4, and *minQ* 20, and multi-allelic SNPs were removed with GATK. The remaining SNPs were then subjected to further filtering with the following parameters: *minDP* 10, *minGQ* 30, *minQ* 30, *min-alleles* 2, *max-alleles* 2, and *maf* 0.05 using VCFtools v0.1.16 (Danecek et al., 2011). Only SNPs with a minimum read depth of 10 were retained using a customized Python3 script.

### Haplotype calling

Per GBS or HiPlex locus, haplotypes were called using the SMAP *haplotype-sites* module within the SMAP package v4.4.0. Read-backed haplotyping was conducted based on the combined variation in SMAPs and SNPs in the GBS read data or based on SNPs in the HiPlex read data using the SMAP *haplotype-sites* module with *mapping_orientation* ignore, *partial* exclude, *no_indels*, *min_read_count* 10, *min_distinct_haplotypes* 2, *min_haplotype_frequency* 5, *discrete_calls* dosage, *frequency_interval_bounds* 10 10 90 90, and *dosage_filter* 2.

### Genetic similarity

The genetic similarity between samples within the Discovery (n=218), Validation (n=105), Screening (n=730), and Canephora Panel (n=514) (composition of panel sets see below) was quantified with the SMAP *grm* module within the SMAP package v4.4.0, using the Jaccard genetic similarity coefficient (Jaccard, 1912) that was calculated based on the discrete dosage haplotype calls in polymorphic GBS or 86 high-quality HiPlex loci. SMAP *grm* was run with *locus_completeness* 0.1, *similarity_coefficient* Jaccard, *distance_method* Euclidean, *locus_information_content* Shared, and *partial* FALSE creating a pairwise Jaccard genetic similarity matrix.

Genetic fingerprints with a Jaccard genetic similarity score greater than 0.977 and 0.965, for GBS data and HiPlex data, respectively, were grouped together and technical and biological replicates (*i.e.* clonal material) were defined. Unique genetic fingerprints were defined by a Jaccard genetic similarity lower than 0.977 and 0.965, for GBS data and HiPlex data, respectively, in pairwise comparison to all other genetic fingerprints.

### HiPlex primer design and sequencing

To construct a HiPlex assay, a minimal set of loci that could discriminate all 139 unique genetic fingerprints in the Discovery Panel was selected according to the following strategy. First, SMAP *delineate* was run on the GBS read data with the following parameters; *mapping_orientation* ignore, *min_stack_depth* 4, *max_stack_depth* 400, *min_cluster_depth* 8, *max_cluster_depth* 400, *max_smap_number* 2, and *completeness* 90. Only loci with a length of 100 – 140 bp were retained. To check for discriminative power across the 218 samples of the Discovery Panel, haplotypes were called for the loci with 2 SMAPs and length 100-140 bp and Jaccard genetic similarities were calculated using the SMAP *grm* module. Second, Primer3 v2.4.0 (Untergasser et al., 2012) was run on the selected loci, taking all known GBS SNP positions into account. Genotype calls within the designed amplicons were simulated using the GBS read data and the amplified loci nested within the GBS loci. To check if each GBS-based genetic fingerprint (based on GBS markers) could still be differentiated using the simulated HiPlex genotype calls, haplotypes were called and Jaccard genetic similarity was calculated using the simulated HiPlex genotype calls. Third, 96 loci with two to four haplotypes per locus were selected, without losing the loci that discriminate the highly similar fingerprints. The primers that amplified these loci were used for HiPlex sequencing to genotype the Yangambi coffee collection samples (n=730; ‘Screening Panel’).

To investigate which genetic resources are present in the Yangambi coffee collection, genomic DNA of all 730 samples collected from the INERA Coffee Collection in Yangambi, was submitted for HiPlex sequencing. HiPlex amplification reactions and library preparations were done by Floodlight Genomics LLC (Knoxville, USA). The libraries were sequenced with 150 PE on a HiSeq3000 instrument (Admera Health, South Plainfield, USA). Forward and reverse reads were merged with PEAR v0.9.11 (Zhang et al., 2014), and the merged reads were aligned to the *C. canephora* reference genome sequence (Denoeud et al., 2014) with the BWA-mem algorithm in BWA v0.7.17 (Li, 2013) with default parameters. Alignments were sorted, indexed, and filtered on mapping quality above 20 with SAMtools v1.10 (Li et al., 2009).

### Validation of the HiPlex assay

To validate the HiPlex assay, we selected samples with minimum 250.000 reads per library. Distribution of reads depth across the 96 HiPlex loci was analyzed to select high-quality HiPlex loci. Ten loci showed completeness lower than 10% and were excluded from further analysis.

We selected samples with sufficient read depth per locus across 86 high-quality HiPlex loci, in both GBS read data and HiPlex read data, further referred to as the ‘Validation Panel’ (n=105) (**Fig 1**), and ran GATK on all these bam files together. To compare the polymorphisms within the GBS and HiPlex SNP data, SNPs within the 86 high-quality HiPlex loci were called with GATK with the same parameters as above (SNP calling) on all GBS and HiPlex bam files, resulting in a SNP call file for the two genotyping techniques together. GBS SNPs were separated from HiPlex using the *keep* function in VCFtools and only polymorphic SNPs were retained by using a minor allele count of 2 for GBS and HiPlex data separately. On a per-sample basis, all SNP genotype calls derived from GBS data were compared to those of HiPlex data across the 86 high-quality HiPlex loci. Read-backed haplotyping was conducted with the SMAP *haplotype-sites* module based on the SNPs within the 86 high-quality HiPlex loci for both GBS and HiPlex read data. On a per-sample basis, all haplotype calls derived from GBS data were compared to those of HiPlex data across the 86 high-quality HiPlex loci (**Fig 2A**).

**Figure 1:**
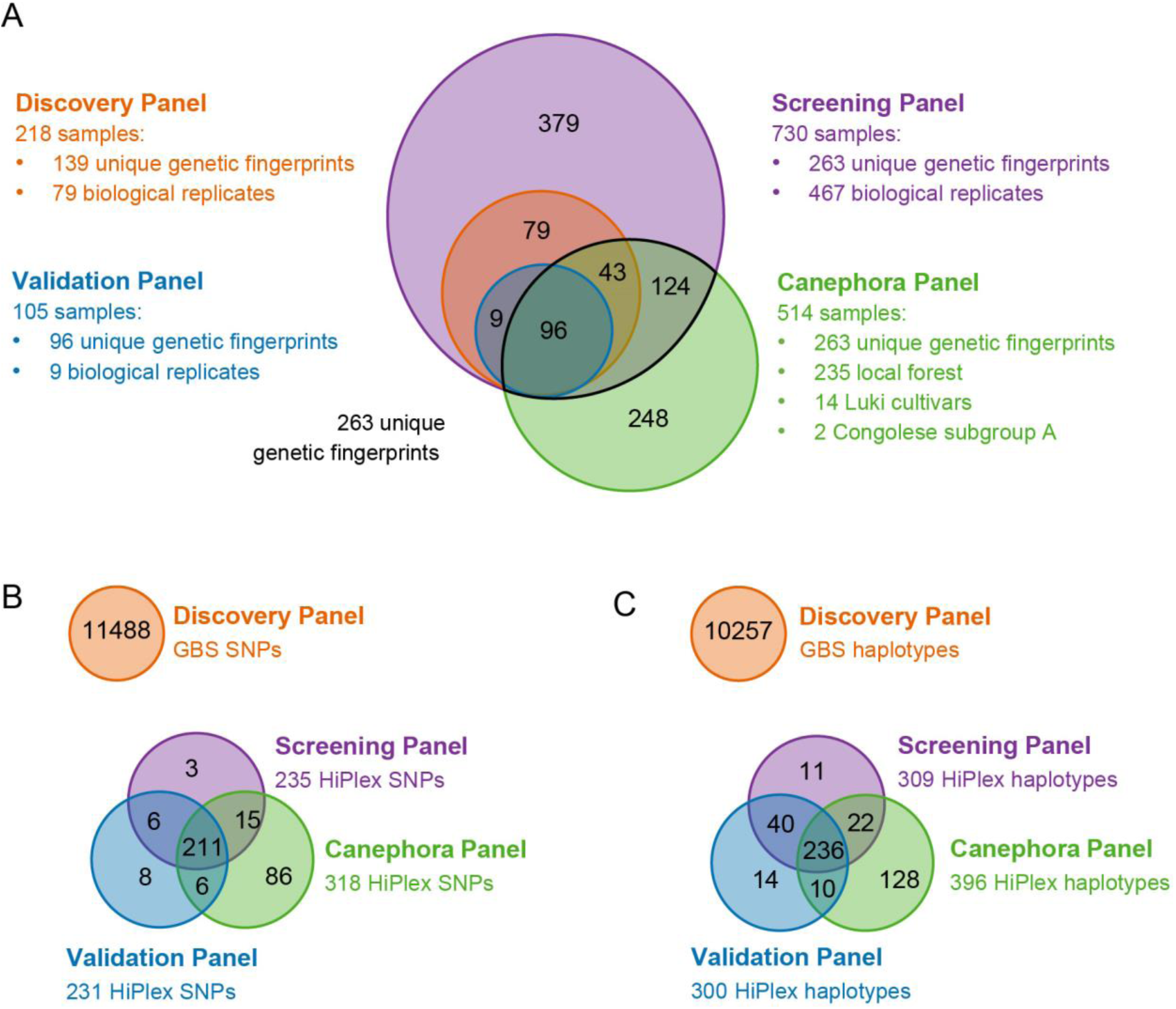
Overview of the Discovery, Validation, Screening, and Canephora Panel set. **(A)** Number of samples used within and shared between the four different panel sets. The black line indicates the unique genetic fingerprints identified within the Yangambi coffee collection. (**B)** Number of unique and common HiPlex SNPs found within and between the Validation, Screening, and Canephora Panel set. The number of GBS SNPs within the Discovery Panel was placed separately to indicate the difference in magnitude between GBS and HiPlex. (**C)** Number of unique and common HiPlex haplotypes found within and shared between the Validation, Screening, and Canephora Panel set. The number of GBS haplotypes within the Discovery panel was placed separately to indicate the difference in magnitude between GBS and HiPlex.

**Figure 2:**
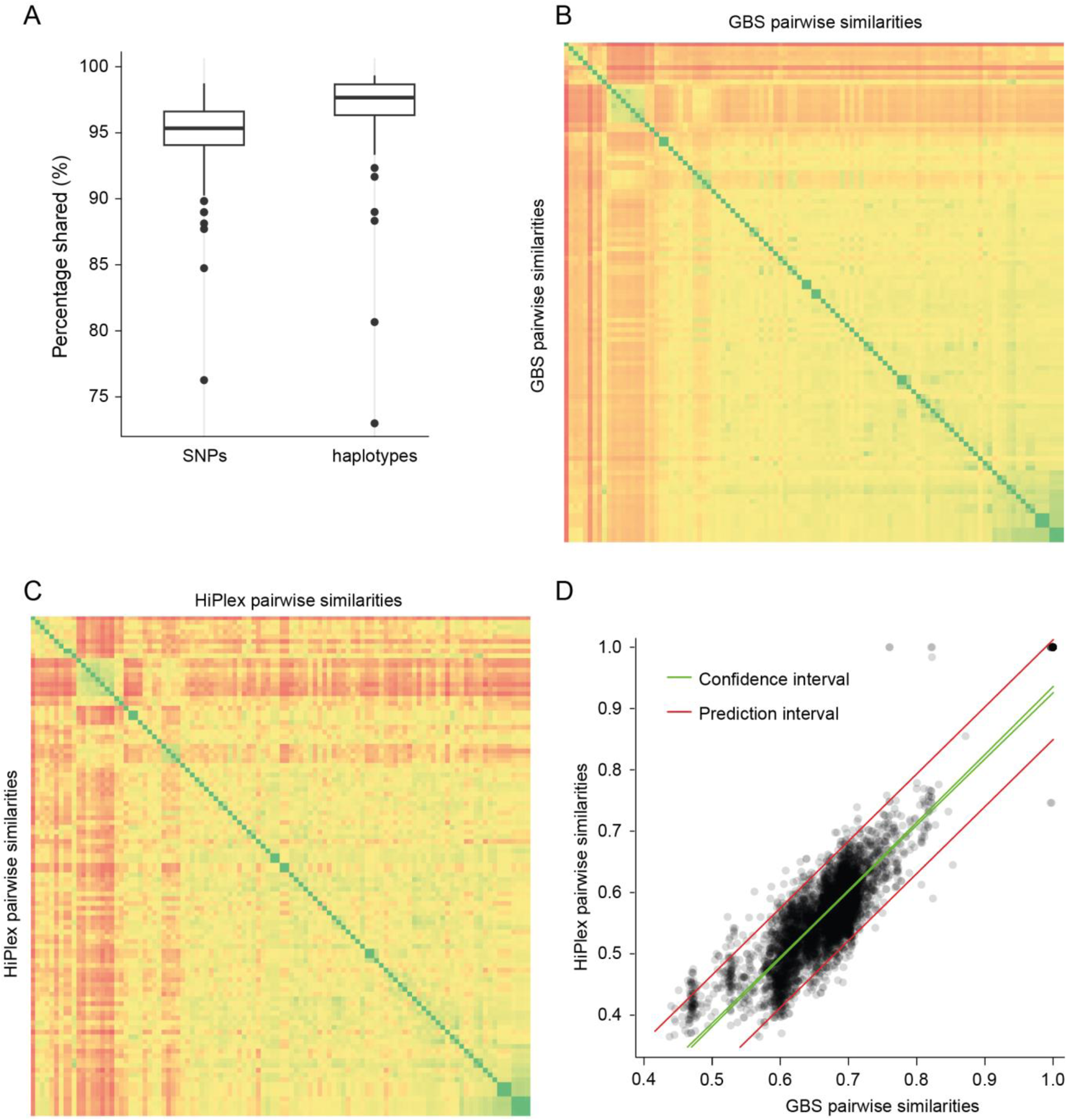
Validation of the HiPlex screening assay. **(A)** Comparison of SNPs and haplotype calls between HiPlex and GBS data. **(B)** Pairwise Jaccard genetic similarity was calculated for all sample pairs within 105 samples based on 10.257 GBS haplotypes within 3125 polymorphic high-quality GBS loci. **(C)** Pairwise Jaccard genetic similarity was calculated for all 105 samples based on 300 HiPlex haplotypes within 86 high-quality HiPlex loci. **(D)** Correlation between Jaccard genetic similarities of GBS (X-axis) and HiPlex (Y-axis), with prediction interval shown as red lines and the 95% confidence intervals as green lines.

Since the selection of HiPlex loci can introduce bias in the calculation of genetic distances between all pairs of samples and the population structure is calculated on these genetic distances, we calibrated whether pairwise Jaccard genetic similarity calculated on GBS data (**Fig 2B**) was correlated to that based on HiPlex data (**Fig 2C**). Pairwise Jaccard genetic similarity was calculated with the same parameters as above (see Genetic similarity), once based on the GBS data within all polymorphic high-quality GBS loci and once based on the HiPlex data within the 86 high-quality HiPlex loci. A correlation between the pairwise Jaccard genetic similarity in the GBS data and HiPlex data was calculated using the Kendall rank correlation test in the ‘ggpubr’ R package (Kassambara & Kassambara, 2020). Prediction values and confidence intervals were analyzed with the *predict* function in R.

### Genetic structure within the Yangambi coffee collection

HiPlex read data of representatives of the 263 unique genetic fingerprints, 14 ‘Luki’ cultivars, 235 wild coffee shrubs, and WGS read data of two samples of the Congolese subgroup A was mapped on the reference genome sequence as described above and used for joined SNP calling on 86 high-quality HiPlex loci. Based on documentation present in the INERA Coffee Collection in Yangambi, seven samples (G0094FOG_2065, G0119FOG_2624, G0105FOG_2676, G0198FOG_2800, G0121FOG_2824, G0138FOG_3231 and G0197FOG_3232) were ‘Lula’ cultivars (Supplemental Table S1). A principal component analysis (PCA) was performed using the R package ADEGENET (Jombart, 2008). Additionally, a Bayesian clustering implemented in fastSTRUCTURE v1.0 (Raj et al., 2014) was run given the most optimal number of genetic clusters (K). Hundred iterations were run for each expected cluster setting K, ranging from 2 to 9. The StructureSelector software (Li & Liu, 2018) was used to determine the most optimal number of K, by first plotting the mean log probability of each successive K and then using the Delta K method following Evanno et al. (2005).

### Parentage analysis

To reveal parent-progeny pairs in the Yangambi coffee collection, a parentage analysis was performed on the 263 unique genetic fingerprints of the collection. We first ran an allele frequency analysis on the 86 high-quality HiPlex loci using the program CERVUS v3.0.7 (Kalinowski et al., 2007). Next, a parentage analysis simulation, which uses a pair-wise likelihood comparison-based approach to assign parent pairs with unknown sexes, was run for 10.000 progenies produced by 263 candidate parents, with 30% parent samples, 50% proportion of loci sampled, 1% proportion of loci mistyped, and confidence levels assessed by LOD distribution (relaxed >80%, strict >95%). Based on the simulation, all 263 unique genetic fingerprints were tested as both progeny and parent, and only parent-progeny pairs with strict (>95%) confidence levels were retained.

### Establishment of a core collection

Core Hunter 3 v3.2.0 R package (De Beukelaer et al., 2018) was used to test two core collection strategies for the 263 unique genetic fingerprints within the Yangambi coffee collection: the M strategy (hereafter referred to as CC-I), which focuses on selecting the most diverse loci to maximize the genetic diversity and; the genetic distance method (hereafter referred to as CC-X), which aims to select the most diverse plant material within a collection to maximize the genetic distance between the entries of the core collection. For both core types allele coverage (CV), diversity within and between alleles (*He* and Shannon’s index (SH)), and average genetic Modified Roger’s (MR) distance between entry-accession (AN) and entry-to-nearest-entry (EN) were calculated for nine different core sizes (3, 5, 10, 25, 50, 100, 150, 200 and 263 accessions). Core collection size 263 is a representation of all unique genetic fingerprints currently identified in the Yangambi coffee collection. For CC-I, an optimal core size was determined based on maximized genetic diversity (He and SH) and minimized AN distance. For CC-X, an optimal core size was determined based on maximized genetic diversity (*He* and SH) and maximized EN distance. The function *set.seed* 100 was used to eliminate randomness in assigning accessions to the core subsets. Accessions were assigned to the core subset using the function *sampleCore* within the Core Hunter 3 R package with *objective* AN (CC-I) or EN (CC-X) and MR, *steps* 500, and *size* equals the optimal core size.

## RESULTS

### Discovery of the genetic diversity in the Yangambi coffee collection

Based on available documentation (field maps and plant labels), a set of 218 samples (Discovery Panel) was selected from the INERA Coffee Collection in Yangambi and subjected to GBS, yielding 7627 high-quality GBS loci with read depth >8 in more than 90% of the samples. These GBS loci had two types of polymorphic sites, namely 18.225 read mapping polymorphisms (SMAPs) identified by the SMAP *delineate* module and 11.488 SNPs called with GATK (referred to as GBS SNPs; **Fig 1**). The polymorphic sites (SNPs and SMAPs) were converted into 10.257 haplotypes using the SMAP *haplotype-sites* module (referred to as GBS haplotypes), yielding a genome-wide marker set of 3177 polymorphic high-quality GBS loci.

The genetic similarity matrix, constructed with the SMAP *grm* module, was arranged to reveal blocks of sample pairs with high similarity (**Fig S1**). In addition, the distribution of all pairwise Jaccard genetic similarity values showed a group of sample pairs with similarities greater than 0.977, which, as expected, contained known replicates (**Fig S1**). Therefore, we set a threshold of minimal Jaccard genetic similarity of 0.977 to identify all pairs of genetically identical samples (*i.e.* clones). In turn, this threshold was used to group all samples into a minimal set of unique genetic fingerprints, yielding 139 groups with unique genetic fingerprints (here labeled like G0001) (**Table S1).** Pairwise Jaccard genetic similarity between the 139 unique genetic fingerprints ranged between 0.44 and 0.89. This grouping confirmed that *a priori* known replicates were genetically identical. However, this analysis also revealed that within unique genetic groups, some individuals had distinct ‘documented identities’ according to field maps and plant labels, indicating incorrect labeling. Conversely, some individuals with the same documented identity were divided over distinct genetic groups, *e.g.* samples with documented identity ‘YB001’ comprised a total of four different unique genetic fingerprints with a Jaccard similarity range of 0.67 to 0.81 (**Table S1**).

### Development of a HiPlex screening assay to routinely screen the collection

First, 483 GBS loci without read mapping polymorphisms (*i.e.* two SMAPs) and locus length between 100 and 140 bp were selected within the 3177 polymorphic high-quality GBS loci set of the Discovery Panel. Within those 483 loci, and taking all known GBS SNP positions into account, Primer3 could successfully design 205 amplicons. Of these 205 loci, 96 loci were selected based on haplotype complexity (range of two to four haplotypes per locus), that together could discriminate all 139 unique genetic fingerprints. This HiPlex assay was then performed on all 730 samples collected from the INERA Coffee Collection in Yangambi, on 235 samples collected from the local rainforest in the Yangambi region (Depecker et al., 2023), and also on 14 ‘Luki’ cultivars supplied by Meise Botanic Garden.

Eighty-six high-quality HiPlex loci gave sufficient read depth across samples with minimum 250.000 reads per library and were retained for further analysis. Hundred-and-five samples with sufficiently deep sequencing per locus in both GBS and HiPlex data were used as Validation Panel. This panel (**Fig 1**) yielded 235 SNPs in the entire set, of which 213 polymorphic SNPs were found for GBS, 231 polymorphic SNPs for HiPlex, and 210 SNPs were found in both. The 235 SNPs were converted into 300 haplotypes within 86 loci.

On a per-sample basis, all genotype calls of GBS data were compared to those of HiPlex data across the 86 loci, yielding a per-sample genotype call reproducibility across the two genotyping techniques of, on average, 94.8% +/- 3.18 (SD) across all samples (**Fig 2A**). On a per-sample basis, all haplotype calls of GBS data were compared to those of HiPlex data across the 86 loci, yielding a per-sample haplotype call reproducibility across the two genotyping techniques of 96.8% +/- 3.52 (SD) across all samples (**Fig 2A**).

Jaccard genetic similarity was calculated on GBS data (**Fig 2B**), of which 18.225 SMAPs and 11.488 SNPs were converted into 10.257 haplotypes within 3125 loci, and on HiPlex data (**Fig 2C**), of which 231 SNPs were converted into 300 haplotypes within 86 loci. The pairwise Jaccard genetic similarity of the GBS data showed a positive correlation with the pairwise Jaccard genetic similarity of the HiPlex data in corresponding sample pairs (R^2^ Adj.= 0.78, p <0.05) showing that the genetic similarities estimated by the genome wide markers (GBS) are accurately reflected by the much smaller set of 86 HiPlex markers (**Fig 2D**).

Next, HiPlex data was analyzed across 730 samples of the Screening Panel, yielding 235 SNPs (screening SNPs) and 309 haplotypes (screening haplotypes) within 86 loci (**Fig 1**). Pairwise Jaccard genetic similarity was calculated on the 309 haplotypes and the distribution of all pairwise similarity values showed a group of replicates with similarities greater than 0.965. Using the minimal genetic similarity of 0.965 as a threshold to collapse individuals into groups of individuals with shared unique genetic fingerprints, the 139 unique genetic fingerprints, initially identified within the Discovery Panel, were reconstituted and complemented by 124 novel genetic fingerprints, resulting in a total of 263 unique genetic fingerprints (further referred to as G0001-G0263) (**Table S1**). Pairwise Jaccard genetic similarity between the 263 unique genetic fingerprints based on the screening haplotypes ranged between 0.32 and 0.88.

### Genetic structure of the Yangambi coffee collection

HiPlex data of 14 ‘Luki’ cultivars, 235 wild coffee shrubs, and 263 unique genetic fingerprints within the Yangambi coffee collection and WGS data of two representatives of Congolese subgroup A (‘Canephora Panel’; n=514) was analyzed yielding a total of 318 SNPs and 396 haplotypes within 86 high-quality HiPlex loci (**Fig 1**). Pairwise Jaccard genetic similarity was calculated on the 396 haplotypes and the distribution of all pairwise similarity values showed a group of replicates with similarities greater than 0.977. Using the minimal genetic similarity of 0.977 as a threshold, one replicate was found for the ‘Luki’ cultivars and three replicates were found for the wild samples resulting in 510 unique genetic fingerprints identified. All 514 samples of the Canephora Panel were then used to further explore the genetic structure in the Yangambi coffee collection and determine the presence of these genetic resources.

The PCA performed on 318 SNPs of the Canephora Panel showed that all 235 samples collected from the local rainforest of the Yangambi region (Depecker et al., 2023) clustered together on the positive PC1-axis (**Fig 3A**). ‘Luki’ cultivars clustered together with the two Congolese subgroup A samples on the positive PC2-axis, and ‘Lula’ cultivars on the negative PC1 and PC2-axes. Unique genetic fingerprints, identified within the Yangambi coffee collection, were scattered between these three clusters. Second, Bayesian clustering, implemented in fastSTRUCTURE, was performed on 318 SNPs of the Canephora Panel, revealing three genetic clusters (**Fig 3B**). Similar to the PCA, the cluster analysis separated wild coffee shrubs collected from the local rainforest in the Yangambi region, hereafter referred as wild samples, from Congolese subgroup A and from ‘Lula’ cultivars. All three genetic clusters were present in the collection, including 181 samples with ‘Lula’ ancestry (Q >80%), nine with wild ancestry, and four with Congolese subgroup A ancestry. Twenty-nine samples from the collection that were located between the ‘Lula’ cultivars and wild samples in the PCA showed an admixed ancestry with partial ‘Lula’ and wild ancestry. Thirty-two Yangambi coffee collection samples positioned close to the Congolese subgroup A in the PCA showed an admixed ancestry proportion of ‘Lula’ and Congolese subgroup A ancestry. All 14 samples collected from the INERA Coffee Collection in Luki were assigned to the Congolese subgroup A. The parentage analysis performed on the 263 unique genetic fingerprints with 227 SNPs and 299 haplotypes within the 86 high-quality HiPlex loci assigned 126 progenies and 108 parents in total, of which 39 samples were identified only as progenies, 21 only as parent, and 87 as both progenies and parent revealing a complex network of hybridization (**Fig 3C**). Only eight unique genetic fingerprints were often identified as parent (**Fig 3A, C**). Based on the fastSTRUCTURE results and parentage analysis, 75 progenies were assigned to ‘Lula’, four to Congolese subgroup A, nine to wild, 24 to ‘Lula’-subgroup A hybrid, and 14 to ‘Lula’-wild hybrid. For the parents, 58 unique genetic fingerprints were assigned to ‘Lula’, four to Congolese subgroup A, nine to wild, 21 to ‘Lula’-subgroup A hybrid, and 16 to ‘Lula’-wild hybrid.

**Figure 3:**
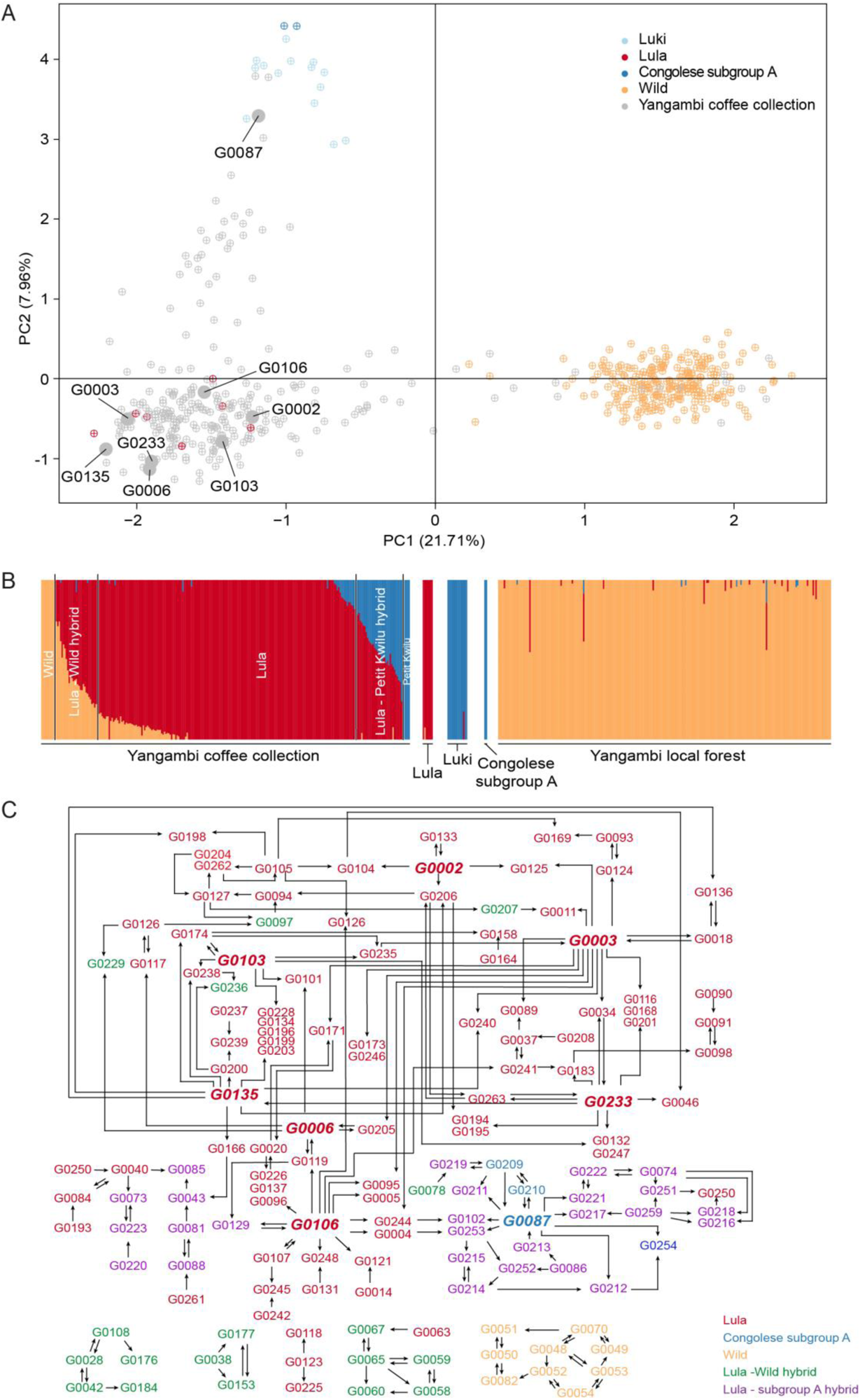
Population genetic structure and parent-progeny pairs within the Yangambi coffee collection. **(A)** Principal component analysis of the Canephora Panel using 318 SNPs indicating ‘Lula’, ‘Luki’, Congolese subgroup A, and wild genetic resources. G-codes indicate the unique genetic fingerprints which were more than five times identified as a parent. **(B)** fastSTRUCTURE bar plot representing three clusters (K = 3). Colors define subpopulations: green (wild), blue (Congolese subgroup A), and red (‘Lula’ cultivars). **(C)** Parentage analysis was performed on the 263 unique genetic fingerprints showing parent-progeny pairs with confidence levels higher than 95%. The arrow indicates which sample is the parent of which progeny.

Comparison of HiPlex SNPs and haplotypes of the Canephora Panel to the HiPlex SNPs and haplotypes of the Discovery, Validation, and Screening Panel, revealed 86 SNPs and 258 haplotypes that were unique to the Canephora Panel (**Fig 1B, C**), showing that the HiPlex assay is able to detect novel SNPs and haplotypes.

### Establishment of a core collection

Two core collection strategies, CC-I and CC-X, were tested for the 263 unique genetic fingerprints within the Yangambi coffee collection. For CC-I, the optimal core size (see Materials and Methods) comprised 100 unique genetic fingerprints as the genetic diversity was higher than other core sizes (*He* of 0.20 and SH of 5.75), all alleles were accounted for (CV of 1) and entry-to-accession distance was low (AN of 0.13) (**Fig 4A**). The accessions assigned to the CC-I core collection were evenly distributed across the ordination space of the Yangambi coffee collection (**Fig 4D**). For CC-X, the optimal core size comprised 10 unique genetic fingerprints as the genetic diversity (*He* of 0.23 and SH of 5.79) was higher than other core sizes tested for CC-X, almost all alleles were accounted for (CV of 0.93) and entry-to-nearest-entry distance was high (EN = 0.35) (**Fig 4B**). The accessions assigned to the CC-X core collection consisted of one Congolese subgroup A genotype, one wild genotype, five ‘Lula’ cultivars, one ‘Lula’-subgroup A hybrid, and two ‘Lula’-wild hybrids **(Fig 4E**).

**Figure 4:**
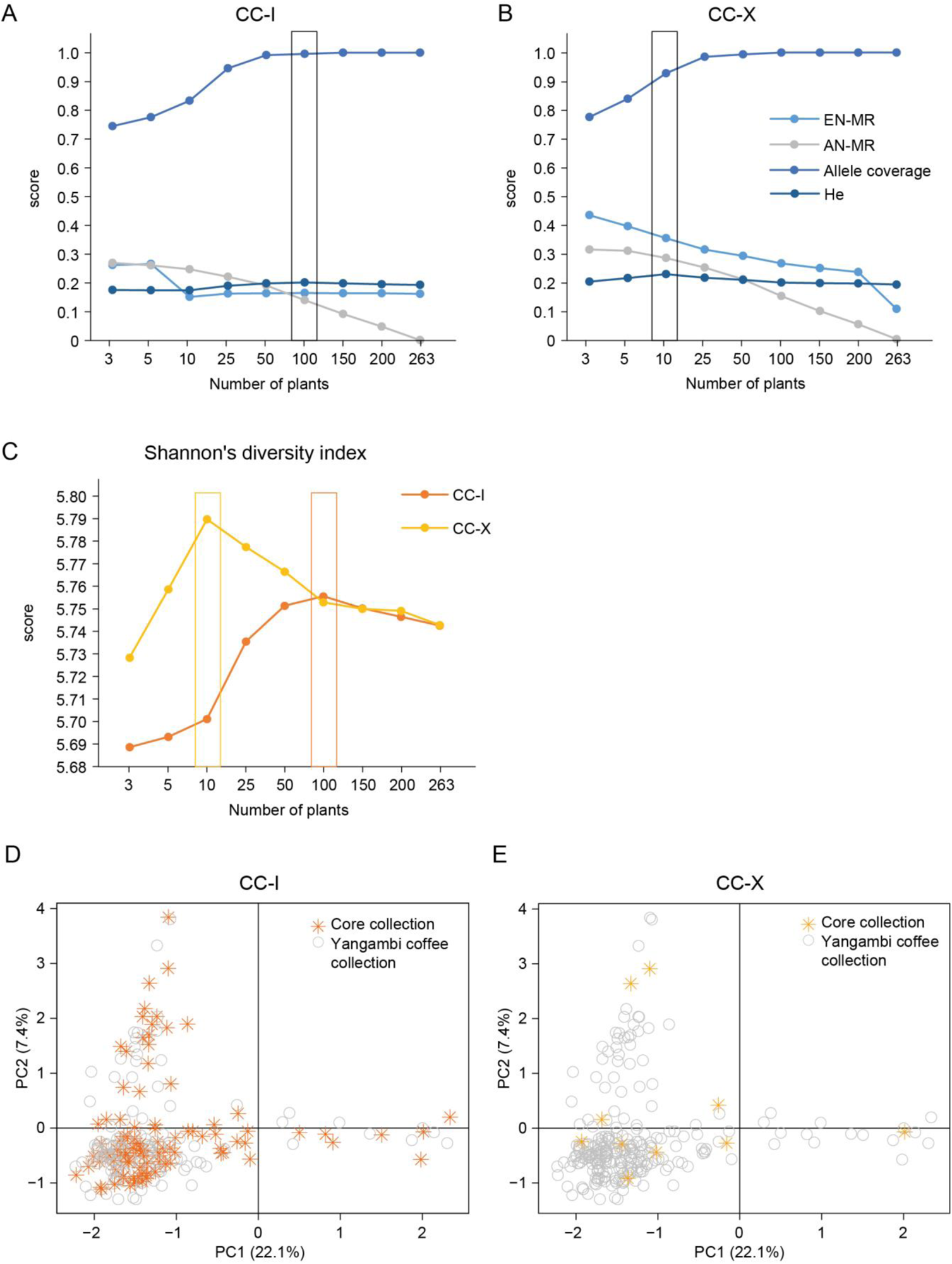
Simulation of different core sizes for core collection strategies CC-I and CC-X. **(A)** Genetic diversity (Allele coverage and He) and distances measures (AN and EN) for core collection strategies CC-I. **(B)** Genetic diversity (Allele coverage and He) and distances measures (AN and EN) for core collection strategies CC-X. **(C)** Shannon’s index was calculated for CC-I and CC-X. Core collection size 263 is a representation of all unique genetic fingerprints currently identified in the Yangambi coffee collection. A principal component analysis was performed on the 263 unique genetic fingerprints within the Yangambi coffee collection indicating the assigned accessions of core collection CC-I **(D)** and CC-X **(E)**.

## DISCUSSION

To enable sustainable conservation and future use of the historically important *C. canephora* collection in Yangambi (DRC), the characterization of its genetic composition is critical. Here, we complemented genome-wide GBS marker sets with a novel versatile HiPlex assay, allowing genetic screening of the INERA Coffee Collection in Yangambi and creating opportunities for future genetic screening in a cost-effective manner. By using the HiPlex screening assay, we were able to reveal the genetic structure and the relative contribution of clonal propagation and crossing products in the collection. In addition, we proposed two core collections which could facilitate the *ex-situ* conservation of *C. canephora* genetic resources of the INERA Coffee Collection in Yangambi in the future.

### Reestablishment of plant labeling

A previous investigation of the INERA Coffee Collection management revealed that the field maps and plant labels of the collection did not correspond with the accessions present in the field (based on field observations 2020-2021, data not shown). Additionally, many plant labels were lost for a substantial portion of the collection. Most clonal-propagated accessions do not differ much in terms of morphology from seed-propagated accessions, whereby it is hard to distinguish them by sight in the field. Therefore, genetic screening of the collection would help to distinguish with certainty the clonal-propagated accession from the seed-propagated accessions. In the present study, a total of 730 samples comprising 117 different plant labels (**Table S1**) were genotyped at 86 high-quality HiPlex loci, and a total of 263 unique genetic fingerprints were identified. Our study revealed that the unique genetic fingerprints did not correspond with the plant labels at individual shrubs. For instance, the number of unique genetic fingerprints is more than two times the number of current labels present in the collection, which means that shrubs carrying the same plant label can be divided into distinct genotypes. Conversely, within genetic groups, some individuals carried different plant labels.

Mislabeling accessions is a common problem in *ex-situ* collections or genetic stocks in general (Bergelson et al., 2016). For example, Akpertey et al. (2021) showed that 18.6% of 400 *C. canephora* accessions from a coffee improvement program at the Cocoa Research Institute of Ghana (CRIG) were mislabeled. They suggested that the majority of the mislabeling originated from the nursery, where trees with the same genotype profile got a different label, or from the wrong replacement of dead plants in the field. In our case, we suspect that the labelling on field became inconsistent with the genetic identities as many accessions were poorly documented and missing their original plant label. Apart from accidental mislabeling, the discrepancy between plant labels and the genetic identity may be the result of labeling practice where siblings are labeled according to parental lines while genetic fingerprints discriminate genetically unique siblings resulting from seed propagation of the collection. To re-establish correct plant labeling, all 730 samples currently collected from the INERA Coffee Collection in Yangambi, were assigned a new label (G0001 – G0263) corresponding to their unique genetic fingerprint obtained with the HiPlex assay.

### A novel versatile HiPlex genotyping assay for routine screening

Different types of molecular markers have previously been used to genotype *C. canephora* cultivars (sequencing based markers and their use in the coffee collections were reviewed by Vi et al 2022), including DArTseq markers for cultivated *C. canephora* in Vietnam and Mexico (Garavito et al., 2016), sequence-related amplified polymorphism (SRAP) and start codon targeted (SCoT) markers for a *C. canephora* germplasm collection in India (Huded et al., 2020), SSR and AFLP markers for a *C. canephora* gene pool in India (Prakash et al., 2005), SSR and RFLP for the CNRA collection in Côte-d’Ivoire (Gomez et al. 2009) or only SSR markers for a *C. canephora* germplasm collection in Brazil (Souza et al., 2013) and for the INERA Coffee Collection in Yangambi that was the subject of this study (Vanden Abeele et al., 2021). Here, we opted to use highly multiplex (HiPlex) amplicon sequencing as it is a simple, accurate, and cost-effective amplicon-based targeted DNA sequencing technique (Kumar et al., 2010). We created a validated HiPlex assay that discriminates the unique genetic fingerprints within the Yangambi coffee collection. We choose for the HiPlex method because it can routinely identify genetic polymorphisms at a set of 86 predefined loci, spread across the genome. Using read-backed haplotyping with the SMAP *haplotype-sites* module, we can transform bi-allelic SNP data to multi-allelic haplotypes to increase genetic resolution per locus. In turn, the combination of all haplotypes defines a unique genetic fingerprint. In addition, since HiPlex is based on resequencing, it allows for discovering novel genetic polymorphisms at all three hierarchical levels (SNPs, haplotypes, and unique genetic fingerprints). For example, 235 SNPs (respectively, 309 haplotypes, and 263 unique genetic fingerprints) were identified in 730 samples representing the Yangambi coffee collection, while 86 novel SNPs (respectively, 128 novel haplotypes, and 247 novel unique genetic fingerprints) were identified in the Canephora Panel, which includes 251 novel samples external to the Yangambi coffee collection (**Fig 1**). Identification of new SNPs made it possible to distinguish cultivated from wild samples, and representatives of the Congolese subgroup A from ‘Lula’ cultivars (**Fig 3**). Since the HiPlex assay can detect new SNPs within the targeted loci, the tool can also be used to genotype yet other *C. canephora* collections to obtain a global overview of the genetic diversity of *C. canephora* collections and compare their genotypes with the unique genetic fingerprints of the Yangambi coffee collection.

Furthermore, genetic fingerprints can have the same SNPs and haplotypes but still differ from each other based on their linear combination of SNPs and haplotypes. Therefore, we can still identify novel unique genetic fingerprints based on Jaccard genetic similarity even if the number of SNPs and haplotypes barely differ between sample sets. For example, 96 unique genetic fingerprints were identified in 105 samples of the Validation Panel (respectively 231 SNPs and 300 haplotypes), while 167 novel unique genetic fingerprints were identified in the Screening Panel (respectively three novel SNPs and 11 novel haplotypes), that includes 625 novel samples external to the Validation Panel (**Fig 1**). Therefore, if the remaining coffee shrubs from the pre-2020 INERA Coffee Collection in Yangambi are genotyped, we suspect that the number of SNPs and haplotypes will not increase substantially but the number of unique genetic fingerprints will increase.

Finally, comparing GBS data to HiPlex data illustrates the trade-off between the cost and efficiency of genotyping versus accuracy in the quantitative estimation of genetic similarities and population structure. An attempt to maintain a minimum number of loci that captures global genetic diversity while enforcing the discriminative power between closely related genotypes slightly inflated the genetic distances between highly similar unique genetic fingerprints. This means that the HiPlex assay is ideal for high-throughput screening of diverse genetic materials, to identify replicates and unique genetic fingerprints, but at the cost of a slight bias in the quantitative estimation of genetic relationships. In addition, while the HiPlex assay is designed to maximize the diversity in the Yangambi coffee collection, and is capable to discover new polymorphisms, it still needs to be optimized to capture diversity external to the pre-2020 INERA Coffee Collection in Yangambi. The strategy outlined here combined with the materials described by Gomez et al. 2009, Merot-L’anthoene et al. (2019) at the species native level, Kiwuka et al. (2022) for Uganda, and Vanden Abeele et al. (2021) for the DRC can be used to develop novel primer sets to complement the current HiPlex assay with a more balanced distribution of genetic diversity in *C. canephora* material worldwide.

### Genetic structure and origin of the Yangambi coffee collection

From 1930 to 1960, the INERA Coffee Collection in Yangambi consisted of Robusta derived material from the Java Research Station, INERA Coffee Research Station in Lula and other wild and cultivated material from the DRC (*e.g.*, from the INERA Coffee Collection in Luki), and abroad. Due to the many difficulties the DRC has faced during the last decades, many of these Robusta cultivars and other wild and cultivated material have been lost. To restore the INERA Coffee Collection in Yangambi, since 2016 the collection is being continuously enriched with numerous new accessions including wild and cultivated material collected mainly by local botanists in several regions within the DRC. To investigate the genetic diversity of the INERA Coffee Collection in Yangambi, the unique genetic fingerprints within the Yangambi coffee collection were compared to ‘Lula’ cultivars, ‘Luki’ cultivars from the INERA Coffee Collection in Luki, reference sample for the Congolese subgroup A (Merot-L’anthoene et al., 2019), and wild samples collected from the local rainforest of the Yangambi region (Depecker et al., 2023). The PCA and structure analyses showed that materials derived from three different genetic resources are present in the INERA Coffee Collection in Yangambi, which is in line with the observations of Vanden Abeele et al. (2021). Most of the unique genetic fingerprints are highly similar to the ‘Lula’ cultivars and some are highly similar to Congolese subgroup A or the local wild genotypes (**Fig 3A**). Vanden Abeele et al. (2021) could identify very few local wild genotypes within the INERA Coffee Collection in Yangambi and therefore proposed to enrich the collection with local wild genotypes to increase the genetic resources available for future breeding and conservation purposes. Our study surveyed a much broader sample set of the INERA Coffee Collection in Yangambi and consequently identified more local wild genotypes (nine in total) than Vanden Abeele et al. (2023). This is consistent with the expansion of the collection with local wild materials as part of the recent rehabilitation initiatives.

In the study of Merot-L’anthoene et al. (2019), wild material that was collected from the same rainforest in the Yangambi region, before the study of Depecker et al. (2023), was assigned to the Congolese subgroup BE (a hybrid between subgroup B and E). Therefore, we can assume that the wild coffee shrubs collected by Depecker et al. (2023) also belong to the Congolese subgroup BE. This ensures that there are currently two different origin groups, namely Congolese subgroup A and BE, present in the collection, but our data shows that the ‘Lula’ cultivars are not assigned to either group. Based on information within the archives of INERA, ‘Lula’ cultivars are assumed to be derived from crossings between ‘*Coffea robusta* L.’ and ‘*Coffea sankuriensis’* but originally collected in the Sankuru region (DRC) (Léonard et al. 2023, in preparation). In order to reveal the origin of the ‘Lula’ cultivars the material have to be compared to each of the eight genetic groups as defined by Merot-L’anthoene et al. (2019). In contrast to the ‘Lula’ cultivars, we were able to assign the ‘Luki’ cultivars to the Congolese subgroup A because they were genetically similar to the corresponding reference material (Merot-L’anthoene et al., 2019).

The Structure analysis showed that around one-quarter of the unique genetic fingerprints had a hybrid identity (29 ‘Lula’-wild hybrid and 32 ‘Lula’-subgroup A hybrid) (**Fig 3B**). These hybrid identities could be a result of dedicated crosses or open pollination. In addition, 29 ‘Lula’-wild hybrid genotypes were found indicating that local wild genotypes are already being used for crossing activities. To investigate the contribution of breeding to the observed genetic structure in the Yangambi coffee collection, a parentage analysis was performed on the 263 genetic fingerprints, which revealed a complex network of hybridization (**Fig 3C**). Most of the parent-progeny pairs found were crosses between ‘Lula’ cultivars and some parent-progeny pairs found were crosses between only wild genotypes within the coffee collection. No specific parent-progeny pairs between ‘Lula’ and Congolese subgroup A, ‘Lula’ and wild or Congolese subgroup A and wild were discovered, but we uncovered multiple parent-progeny pairs between ‘Lula’ and ‘Lula’-subgroup A hybrids. Moreover, eight unique genetic fingerprints were frequently identified as a parent, of which seven were identified as ‘Lula’ cultivars and one as Congolese subgroup A. As Capot (1962) noted that in 1951 seven mother plants were selected for seed distribution, these unique genetic fingerprints may correspond to the ancient selected mother plants or direct descendants thereof.

### Establishment of a core collection using two strategies

There are multiple strategies to construct a core collection, but in recent years, the maximization strategy, which aims to maximize the genetic diversity, and the genetic distance method, which aims to maximize the genetic distance, are the two most commonly used strategies (Gu et al., 2023). In this study, we applied these two core collection strategies to the Yangambi coffee collection, representing the screened genetic diversity within the 263 unique genetic fingerprints (**Fig 4**). We used Core Hunter as this software optimize the genetic distance and allelic diversity simultaneously by weighting the Modified Roger’s distance and Shannon diversity index differently based on entry-accession (AN) and entry-to-nearest-entry (EN) distance (De Beukelaer et al., 2018). The main objective of the maximization strategy (CC-I) was to select the most diverse loci to maximize the genetic diversity and by this maintain a uniform representation of the original genetic diversity. We found that the optimal core size would be 100 entries, capturing all alleles in the Yangambi coffee collection. The main objective of the genetic distance method (CC-X) was to select the most diverse genotypes to maximize the genetic distance between the entries of the core collection and is rather orientated more towards breeding activities. Here, the optimal core collection size of 10 entries captures 93% of all alleles in the Yangambi coffee collection with a maximal entry-to-nearest-entry distance. For genetic resource conservation purposes, the CC-I strategy is the most suited but with an optimal core size of 100 entries, it would be a less cost-effective approach than the CC-X strategy with a core size of 10 individuals. Notably, these core collections were proposed based on genetic diversity without considering the phenotypic or agronomic traits present in the INERA Coffee Collection in Yangambi. If a core collection oriented towards breeding activities would be established for the INERA Coffee Collection, it would be necessary to expand the CC-X core collection with phenotypical or agronomical interesting accessions.

## Conclusion

Following the best practices for validating the identity of genetic stocks (Bergelson et al., 2018), we created a HiPlex assay to routinely check plant labelling within the INERA Coffee Collection in Yangambi. Using the HiPlex screening assay, we identified unique genetic fingerprints and revealed which genetic resources are present within the Yangambi coffee collection. In addition, we proposed two core collections that can contribute to the breeding activities and sustainable and effective management of the Yangambi coffee collection. By creating a novel versatile HiPlex screening assay and presenting two core collection strategies for future use, this study is a first step toward the evaluation and contribution to the conservation of coffee genetic resources in the Democratic Republic of the Congo. This HiPlex screening assay can be used to screen other coffee collections (*e.g.* in Ivory Coast and Uganda) and compare them with the INERA Coffee Collection in Yangambi, to better understand of what part of the genetic diversity space of the cultivated *C. canephora* is already captured in the INERA Coffee Collection and which cultivated material should be used to further enrich the collection.

## Author Contributions

TR, OH, PS, RB, and LV designed this study. RB, JK, TS, and BK participated in fieldwork. RB and PS supplied the genomic DNA extracts of the 14 herbarium samples collected from the INERA Coffee Collection in Luki. JD collected the leaf material of 235 wild coffee shrubs from the Yangambi region. VP supplied the WGS data of the two Congolese subgroup A samples. LV executed the lab work. LV, RB, and TR analyzed the data. LV, RB, PS, OH, and TR wrote the manuscript. All authors contributed to finalizing the manuscript.

## Supporting information

Supplemental Figure S1: Pairwise Jaccard genetic similarity calculated for all 218 samples of the Discovery Panel.

Supplemental Table S1:Overview of the unique genetic fingerprints and plant labels for all 730 samples collected from the INERA Coffee Collection

## Acknowledgments

We would like to thank the Institut National pour l’Étude et la Recherche Agronomiques (INERA) for giving access to their collection; Rachel Ndezu and CIFOR (through the FORETS project, funded by the EU 11th Development Fund) for the logistic and administrative support during fieldwork in the Democratic Republic of the Congo. We would also like to express our sincere gratitude to the Ministère de L’Environnement et Développement Durable (MEDD) for their help with obtaining permits (N°008/ANCCB-RDC/SG-EDD/BTB/11/2020, N°004/ANCCB-RDC/SG-EDD/BTB/2021, N°014/ANCCB-RDC/SG-EDD/BTB/11/2021 and N°025/ANCCB-RDC/SG-EDD/BTB/11/2022).

## Data Availability Statement

The data that support the findings of this study are available on request. In accordance with the Democratic Republic of the Congo and international regulations, restrictions apply on the availability of these data, which were used under license for this study.

