## Supplementary figures and images for "Characterization of the genetic composition and establishment of a core collection for the INERA Robusta coffee (*Coffea canephora*) field genebank from the Democratic Republic of the Congo"

### Supplemental Figure S1: Pairwise Jaccard genetic similarity calculated for all 218 samples of the Discovery Panel.

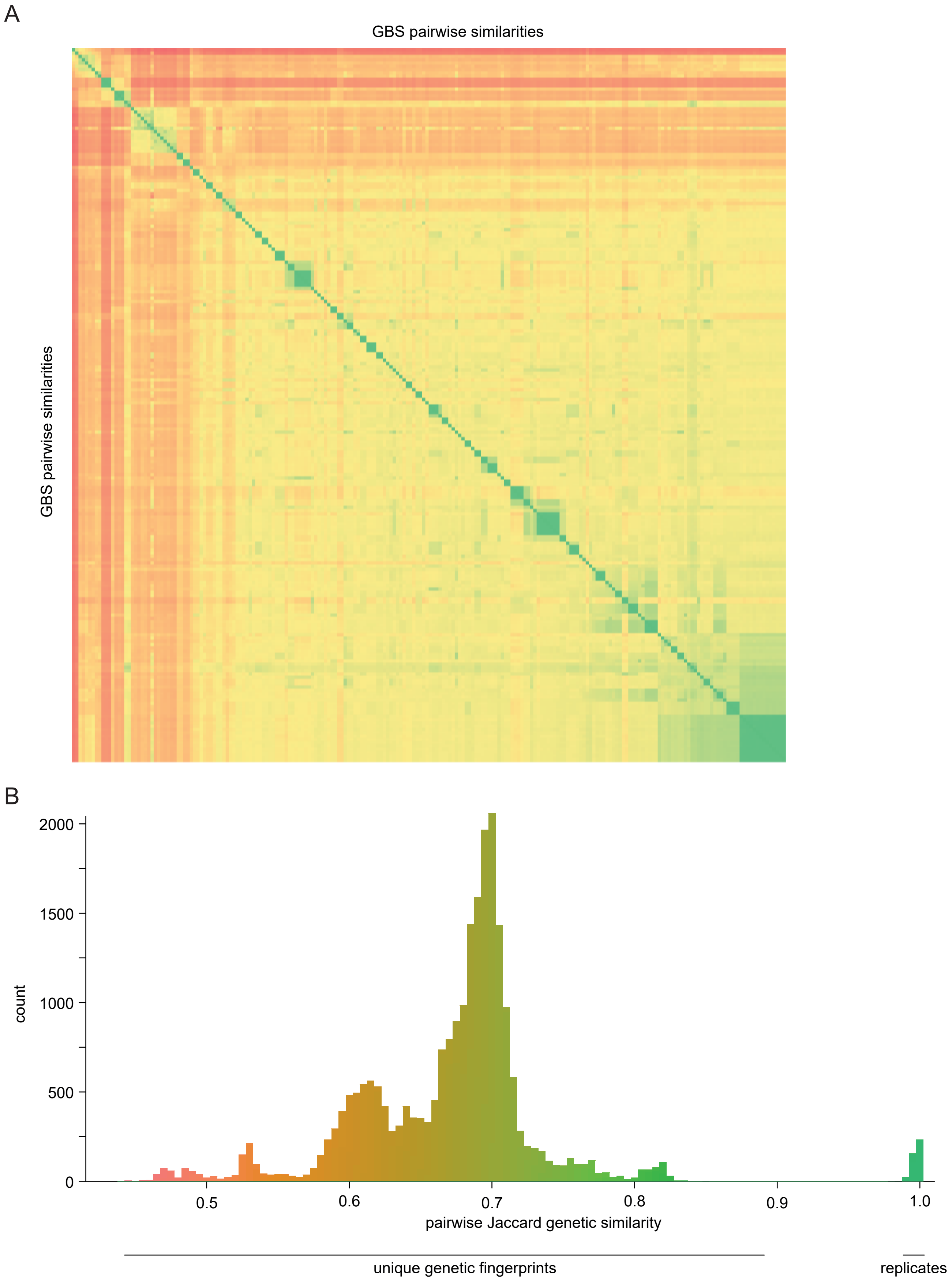
