## Supplemental Table S1:Overview of the unique genetic fingerprints and plant labels for all 730 samples collected from the INERA Coffee Collection for "Characterization of the genetic composition and establishment of a core collection for the INERA Robusta coffee (*Coffea canephora*) field genebank from the Democratic Republic of the Congo"

| Unique sample name | Unique genetic fingerprint | Documented identity | Assigned Genotype | Discovery Panel | Validation Panel | Screening Panel | Canephora Panel | CC-I | CC-X |
|---|---|---|---|---|---|---|---|---|---|
| Collection_G0001FOG_2464 | G0001 | YB0001 | Lula | x | | x | x | | |
| Collection_G0001_2865 | G0001 | YB0001 | Lula | | | x | | | |
| Collection_G0001_2251 | G0001 | YB0001 | Lula | x | x | x | | | |
| Collection_G0001_2875 | G0001 | YB0001 | Lula | | | x | | | |
| Collection_G0001_3008 | G0001 | YB0007 | Lula | | | x | | | |
| Collection_G0001_2344 | G0001 | YB0003 | Lula | | | x | | | |
| Collection_G0002FOG_2608 | G0002 | YB0002 | Lula | x | | x | x | x | |
| Collection_G0002_2667 | G0002 | YB0002 | Lula | | | x | | | |
| Collection_G0002_2668 | G0002 | YB0002 | Lula | | | x | | | |
| Collection_G0002_2804 | G0002 | YB0002 | Lula | | | x | | | |
| Collection_G0002_2890 | G0002 | YB0003 | Lula | | | x | | | |
| Collection_G0002_1966 | G0002 | YB0002 | Lula | x | x | x | | | |
| Collection_G0002_2915 | G0002 | YB0007 | Lula | | | x | | | |
| Collection_G0002_2934 | G0002 | YB0002 | Lula | | | x | | | |
| Collection_G0002_2935 | G0002 | YB0002 | Lula | | | x | | | |
| Collection_G0002_2943 | G0002 | YB0002 | Lula | | | x | | | |
| Collection_G0002_2946 | G0002 | YB0002 | Lula | | | x | | | |
| Collection_G0002_2351 | G0002 | YB0002 | Lula | | | x | | | |
| Collection_G0002_3080 | G0002 | YB0004 | Lula | | | x | | | |
| Collection_G0002_3081 | G0002 | YB0004 | Lula | | | x | | | |
| Collection_G0002_3082 | G0002 | YB0004 | Lula | | | x | | | |
| Collection_G0002_3087 | G0002 | YB0004 | Lula | | | x | | | |
| Collection_G0002_3100 | G0002 | YB0002 | Lula | | | x | | | |
| Collection_G0002_3101 | G0002 | YB0002 | Lula | | | x | | | |
| Collection_G0002_3103 | G0002 | YB0002 | Lula | | | x | | | |
| Collection_G0002_3105 | G0002 | YB0002 | Lula | | | x | | | |
| Collection_G0002_3107 | G0002 | YB0002 | Lula | | | x | | | |
| Collection_G0002_3111 | G0002 | YB0002 | Lula | | | x | | | |
| Collection_G0002_3112 | G0002 | YB0002 | Lula | | | x | | | |
| Collection_G0002_3114 | G0002 | YB0002 | Lula | | | x | | | |
| Collection_G0002_3116 | G0002 | YB0002 | Lula | | | x | | | |
| Collection_G0002_3117 | G0002 | YB0002 | Lula | | | x | | | |
| Collection_G0002_3123 | G0002 | YB0002 | Lula | | | x | | | |
| Collection_G0002_3125 | G0002 | YB0002 | Lula | | | x | | | |
| Collection_G0002_3126 | G0002 | YB0002 | Lula | | | x | | | |
| Collection_G0002_3127 | G0002 | YB0002 | Lula | | | x | | | |
| Collection_G0002_3128 | G0002 | YB0002 | Lula | | | x | | | |
| Collection_G0002_3129 | G0002 | YB0002 | Lula | | | x | | | |
| Collection_G0002_3269 | G0002 | YB0002 | Lula | x | | x | | | |
| Collection_G0002_3254 | G0002 | YB0002 | Lula | x | | x | | | |
| Collection_G0003FOG_2612 | G0003 | YB0003 | Lula | x | | x | x | | |
| Collection_G0003_2670 | G0003 | YB0002 | Lula | | | x | | | |
| Collection_G0003_2671 | G0003 | YB0003 | Lula | | | x | | | |
| Collection_G0003_2672 | G0003 | YB0003 | Lula | | | x | | | |
| Collection_G0003_2808 | G0003 | YB0003 | Lula | | | x | | | |
| Collection_G0003_2850 | G0003 | YB0094 | Lula | | | x | | | |
| Collection_G0003_2876 | G0003 | YB0003 | Lula | | | x | | | |
| Collection_G0003_2877 | G0003 | YB0003 | Lula | | | x | | | |
| Collection_G0003_2878 | G0003 | YB0003 | Lula | | | x | | | |

| Unique sample name | Unique genetic fingerprint | Documented identity | Assigned Genotype | Discovery Panel | Validation Panel | Screening Panel | Canephora Panel | CC-I | CC-X |
|---|---|---|---|---|---|---|---|---|---|
| Collection_G0003_2879 | G0003 | YB0003 | Lula | | | x | | | |
| Collection_G0003_2880 | G0003 | YB0003 | Lula | | | x | | | |
| Collection_G0003_3219 | G0003 | YB0003 | Lula | | | x | | | |
| Collection_G0003_2468 | G0003 | YB0003 | Lula | | | x | | | |
| Collection_G0003_2883 | G0003 | YB0003 | Lula | | | x | | | |
| Collection_G0003_2884 | G0003 | YB0003 | Lula | | | x | | | |
| Collection_G0003_2885 | G0003 | YB0003 | Lula | | | x | | | |
| Collection_G0003_1967 | G0003 | YB0003 | Lula | x | x | x | | | |
| Collection_G0003_2888 | G0003 | YB0003 | Lula | | | x | | | |
| Collection_G0003_2889 | G0003 | YB0003 | Lula | | | x | | | |
| Collection_G0003_2893 | G0003 | YB0003 | Lula | | | x | | | |
| Collection_G0003_2895 | G0003 | YB0003 | Lula | | | x | | | |
| Collection_G0003_2896 | G0003 | YB0003 | Lula | x | x | x | | | |
| Collection_G0003_2901 | G0003 | YB0007 | Lula | | | x | | | |
| Collection_G0003_2902 | G0003 | YB0007 | Lula | | | x | | | |
| Collection_G0003_2903 | G0003 | YB0007 | Lula | | | x | | | |
| Collection_G0003_2912 | G0003 | YB0007 | Lula | | | x | | | |
| Collection_G0003_2916 | G0003 | YB0007 | Lula | | | x | | | |
| Collection_G0003_3243 | G0003 | YB0006 | Lula | x | | x | | | |
| Collection_G0003_2972 | G0003 | YB0007 | Lula | | | x | | | |
| Collection_G0003_2973 | G0003 | YB0007 | Lula | | | x | | | |
| Collection_G0003_2976 | G0003 | YB0007 | Lula | | | x | | | |
| Collection_G0003_2978 | G0003 | YB0007 | Lula | | | x | | | |
| Collection_G0003_2979 | G0003 | YB0007 | Lula | | | x | | | |
| Collection_G0003_2980 | G0003 | YB0007 | Lula | | | x | | | |
| Collection_G0003_2982 | G0003 | YB0007 | Lula | | | x | | | |
| Collection_G0003_2986 | G0003 | YB0007 | Lula | | | x | | | |
| Collection_G0003_2987 | G0003 | YB0007 | Lula | | | x | | | |
| Collection_G0003_2989 | G0003 | YB0007 | Lula | | | x | | | |
| Collection_G0003_2352 | G0003 | YB0007 | Lula | | | x | | | |
| Collection_G0003_2991 | G0003 | YB0007 | Lula | | | x | | | |
| Collection_G0003_2992 | G0003 | YB0007 | Lula | | | x | | | |
| Collection_G0003_2995 | G0003 | YB0007 | Lula | | | x | | | |
| Collection_G0003_2997 | G0003 | YB0007 | Lula | | | x | | | |
| Collection_G0003_2998 | G0003 | YB0007 | Lula | | | x | | | |
| Collection_G0003_2999 | G0003 | YB0007 | Lula | | | x | | | |
| Collection_G0003_3000 | G0003 | YB0007 | Lula | | | x | | | |
| Collection_G0003_2470 | G0003 | YB0005 | Lula | x | x | x | | | |
| Collection_G0003_3054 | G0003 | YB0005 | Lula | | | x | | | |
| Collection_G0003_3073 | G0003 | YB0004 | Lula | | | x | | | |
| Collection_G0003_3090 | G0003 | YB0004 | Lula | | | x | | | |
| Collection_G0003_3094 | G0003 | YB0004 | Lula | | | x | | | |
| Collection_G0003_3227 | G0003 | YB0007 | Lula | x | | x | | | |
| Collection_G0003_3133 | G0003 | YB0007 | Lula | | | x | | | |
| Collection_G0003_3224 | G0003 | YB0001 | Lula | | | x | | | |
| Collection_G0003_3144 | G0003 | YB0001 | Lula | | | x | | | |
| Collection_G0003_3145 | G0003 | YB0001 | Lula | | | x | | | |
| Collection_G0003_3148 | G0003 | YB0001 | Lula | | | x | | | |
| Collection_G0003_3149 | G0003 | YB0001 | Lula | | | x | | | |

| Unique sample name | Unique genetic fingerprint | Documented identity | Assigned Genotype | Discovery Panel | Validation Panel | Screening Panel | Canephora Panel | CC-I | CC-X |
|---|---|---|---|---|---|---|---|---|---|
| Collection_G0003_3264 | G0003 | YB0003 | Lula | x | | x | | | |
| Collection_G0003_3262 | G0003 | YB0003 | Lula | x | | x | | | |
| Collection_G0003_3237 | G0003 | YB0003 | Lula | x | | x | | | |
| Collection_G0003_3261 | G0003 | YB0003 | Lula | x | | x | | | |
| Collection_G0003_3257 | G0003 | YB0003 | Lula | x | x | x | | | |
| Collection_G0003_3234 | G0003 | YB0003 | Lula | x | | x | | | |
| Collection_G0003_3256 | G0003 | YB0003 | Lula | x | | x | | | |
| Collection_G0003_3251 | G0003 | YB0004 | Lula | x | | x | | | |
| Collection_G0004FOG_1968 | G0004 | YB0004 | Lula | x | x | x | x | | |
| Collection_G0004_2469 | G0004 | YB0004 | Lula | | | x | | | |
| Collection_G0004_3064 | G0004 | YB0004 | Lula | | | x | | | |
| Collection_G0004_3066 | G0004 | YB0004 | Lula | | | x | | | |
| Collection_G0004_3079 | G0004 | YB0004 | Lula | | | x | | | |
| Collection_G0004_3083 | G0004 | YB0004 | Lula | | | x | | | |
| Collection_G0004_3086 | G0004 | YB0004 | Lula | | | x | | | |
| Collection_G0004_3253 | G0004 | YB0003 | Lula | | | x | | | |
| Collection_G0005FOG_2929 | G0005 | YB0006 | Lula | x | | x | x | x | |
| Collection_G0005_3013 | G0005 | YB0005 | Lula | | | x | | | |
| Collection_G0005_3019 | G0005 | YB0005 | Lula | | | x | | | |
| Collection_G0005_3025 | G0005 | YB0005 | Lula | | | x | | | |
| Collection_G0005_3029 | G0005 | YB0005 | Lula | | | x | | | |
| Collection_G0005_3031 | G0005 | YB0005 | Lula | | | x | | | |
| Collection_G0005_3032 | G0005 | YB0005 | Lula | | | x | | | |
| Collection_G0005_3037 | G0005 | YB0005 | Lula | | | x | | | |
| Collection_G0005_3038 | G0005 | YB0005 | Lula | | | x | | | |
| Collection_G0005_3039 | G0005 | YB0005 | Lula | | | x | | | |
| Collection_G0005_3040 | G0005 | YB0005 | Lula | | | x | | | |
| Collection_G0005_3041 | G0005 | YB0005 | Lula | | | x | | | |
| Collection_G0005_3042 | G0005 | YB0005 | Lula | | | x | | | |
| Collection_G0005_3049 | G0005 | YB0005 | Lula | | | x | | | |
| Collection_G0005_3050 | G0005 | YB0005 | Lula | | | x | | | |
| Collection_G0005_3052 | G0005 | YB0005 | Lula | | | x | | | |
| Collection_G0005_3226 | G0005 | YB0005 | Lula | x | | x | | | |
| Collection_G0006FOG_2607 | G0006 | YB0002 | Lula | x | | x | x | | |
| Collection_G0006_2616 | G0006 | YB0004 | Lula | | | x | | | |
| Collection_G0006_2669 | G0006 | YB0002 | Lula | | | x | | | |
| Collection_G0006_2679 | G0006 | YB0004 | Lula | | | x | | | |
| Collection_G0006_2819 | G0006 | YB0006 | Lula | | | x | | | |
| Collection_G0006_2822 | G0006 | YB0006 | Lula | | | x | | | |
| Collection_G0006_2823 | G0006 | YB0006 | Lula | | | x | | | |
| Collection_G0006_2909 | G0006 | YB0007 | Lula | | | x | | | |
| Collection_G0006_2924 | G0006 | YB0006 | Lula | | | x | | | |
| Collection_G0006_2252 | G0006 | YB0006 | Lula | x | x | x | | | |
| Collection_G0006_3004 | G0006 | YB0007 | Lula | | | x | | | |
| Collection_G0006_3006 | G0006 | YB0007 | Lula | | | x | | | |
| Collection_G0006_3057 | G0006 | YB0005 | Lula | | | x | | | |
| Collection_G0006_3078 | G0006 | YB0004 | Lula | | | x | | | |
| Collection_G0006_3240 | G0006 | YB0004 | Lula | x | | x | | | |
| Collection_G0007FOG_2641 | G0007 | YB0011 | Lula | x | | x | x | | |

| Unique sample name | Unique genetic fingerprint | Documented identity | Assigned Genotype | Discovery Panel | Validation Panel | Screening Panel | Canephora Panel | CC-I | CC-X |
|---|---|---|---|---|---|---|---|---|---|
| Collection_G0007_2829 | G0007 | YB0007 | Lula | | | x | | | |
| Collection_G0007_2253 | G0007 | YB0007 | Lula | x | x | x | | | |
| Collection_G0007_2900 | G0007 | YB0007 | Lula | | | x | | | |
| Collection_G0007_2911 | G0007 | YB0007 | Lula | | | x | | | |
| Collection_G0008FOG_1972 | G0008 | YB0008 | Lula | x | x | x | x | x | |
| Collection_G0008_2618 | G0008 | YB0008 | Lula | | | x | | | |
| Collection_G0008_2619 | G0008 | YB0008 | Lula | | | x | | | |
| Collection_G0008_2680 | G0008 | YB0008 | Lula | | | x | | | |
| Collection_G0008_2681 | G0008 | YB0008 | Lula | | | x | | | |
| Collection_G0008_2682 | G0008 | YB0008 | Lula | | | x | | | |
| Collection_G0008_2684 | G0008 | YB0008 | Lula | | | x | | | |
| Collection_G0008_2685 | G0008 | YB0008 | Lula | | | x | | | |
| Collection_G0008_2686 | G0008 | YB0008 | Lula | | | x | | | |
| Collection_G0008_2687 | G0008 | YB0008 | Lula | | | x | | | |
| Collection_G0008_2688 | G0008 | YB0008 | Lula | | | x | | | |
| Collection_G0008_2689 | G0008 | YB0008 | Lula | | | x | | | |
| Collection_G0008_2690 | G0008 | YB0008 | Lula | | | x | | | |
| Collection_G0008_2691 | G0008 | YB0008 | Lula | | | x | | | |
| Collection_G0008_2692 | G0008 | YB0008 | Lula | | | x | | | |
| Collection_G0008_2693 | G0008 | YB0008 | Lula | | | x | | | |
| Collection_G0008_2694 | G0008 | YB0008 | Lula | | | x | | | |
| Collection_G0008_2695 | G0008 | YB0008 | Lula | | | x | | | |
| Collection_G0008_2696 | G0008 | YB0008 | Lula | | | x | | | |
| Collection_G0008_2697 | G0008 | YB0008 | Lula | | | x | | | |
| Collection_G0008_3203 | G0008 | YB0008 | Lula | x | | x | | | |
| Collection_G0008_2448 | G0008 | YB0008 | Lula | | | x | | | |
| Collection_G0008_2449 | G0008 | YB0008 | Lula | | | x | | | |
| Collection_G0008_2450 | G0008 | YB0008 | Lula | | | x | | | |
| Collection_G0008_2759 | G0008 | YB0008 | Lula | | | x | | | |
| Collection_G0008_2451 | G0008 | YB0008 | Lula | | | x | | | |
| Collection_G0008_2452 | G0008 | YB0008 | Lula | | | x | | | |
| Collection_G0008_2453 | G0008 | YB0008 | Lula | | | x | | | |
| Collection_G0009FOG_2254 | G0009 | YB0009 | Lula | x | x | x | x | | |
| Collection_G0009_2633 | G0009 | YB0009 | Lula | x | | x | | | |
| Collection_G0009_2635 | G0009 | YB0009 | Lula | | | x | | | |
| Collection_G0009_3258 | G0009 | YB0009 | Lula | x | | x | | | |
| Collection_G0009_3223 | G0009 | YB0009 | Lula | x | | x | | | |
| Collection_G0010FOG_2636 | G0010 | YB0010 | Lula-subgroup A | x | | x | x | | |
| Collection_G0010_1974 | G0010 | YB0010 | Lula-subgroup A | x | x | x | | | |
| Collection_G0010_2637 | G0010 | YB0010 | Lula-subgroup A | | | x | | | |
| Collection_G0010_2638 | G0010 | YB0010 | Lula-subgroup A | | | x | | | |
| Collection_G0010_2458 | G0010 | YB0046 | Lula-subgroup A | | | x | | | |
| Collection_G0010_3208 | G0010 | YB0046 | Lula-subgroup A | x | | x | | | |
| Collection_G0010_2984 | G0010 | YB0007 | Lula-subgroup A | | | x | | | |
| Collection_G0010_2993 | G0010 | YB0007 | Lula-subgroup A | | | x | | | |
| Collection_G0010_2996 | G0010 | YB0007 | Lula-subgroup A | | | x | | | |
| Collection_G0010_3003 | G0010 | YB0007 | Lula-subgroup A | | | x | | | |
| Collection_G0010_3005 | G0010 | YB0007 | Lula-subgroup A | | | x | | | |
| Collection_G0010_3007 | G0010 | YB0007 | Lula-subgroup A | | | x | | | |

| Unique sample name | Unique genetic fingerprint | Documented identity | Assigned Genotype | Discovery Panel | Validation Panel | Screening Panel | Canephora Panel | CC-I | CC-X |
|---|---|---|---|---|---|---|---|---|---|
| Collection_G0011FOG_1975 | G0011 | YB0011 | Lula-Wild | x | x | x | x | | |
| Collection_G0012FOG_1976 | G0012 | YB0012 | Lula | x | x | x | x | | |
| Collection_G0012_2643 | G0012 | YB0012 | Lula | x | | x | | | |
| Collection_G0012_2644 | G0012 | YB0012 | Lula | | | x | | | |
| Collection_G0013FOG_1978 | G0013 | YB0014 | Lula-Wild | x | x | x | x | | |
| Collection_G0013_2648 | G0013 | YB0014 | Lula-Wild | | | x | | | |
| Collection_G0013_2649 | G0013 | YB0014 | Lula-Wild | | | x | | | |
| Collection_G0013_2650 | G0013 | YB0014 | Lula-Wild | | | x | | | |
| Collection_G0013_2651 | G0013 | YB0014 | Lula-Wild | | | x | | | |
| Collection_G0013_2661 | G0013 | YB0014 | Lula-Wild | | | x | | | |
| Collection_G0013_2442 | G0013 | YB0014 | Lula-Wild | x | | x | | | |
| Collection_G0013_2663 | G0013 | YB0014 | Lula-Wild | | | x | | | |
| Collection_G0013_2664 | G0013 | YB0014 | Lula-Wild | | | x | | | |
| Collection_G0014FOG_1979 | G0014 | YB0015 | Lula | x | x | x | x | x | |
| Collection_G0015FOG_1980 | G0015 | YB0016 | Lula | x | x | x | x | | |
| Collection_G0015_2269 | G0015 | YB0089 | Lula | x | x | x | | | |
| Collection_G0016FOG_1981 | G0016 | YB0017 | Lula-Wild | x | x | x | x | | |
| Collection_G0017FOG_1982 | G0017 | YB0018 | Lula | x | x | x | x | | |
| Collection_G0017_2660 | G0017 | YB0018 | Lula | x | | x | | | |
| Collection_G0018FOG_2071 | G0018 | YB0016 | Lula | x | x | x | x | x | |
| Collection_G0018_1983 | G0018 | YB0019 | Lula | x | x | x | | | |
| Collection_G0018_1990 | G0018 | YB0026 | Lula | x | x | x | | | |
| Collection_G0019FOG_2654 | G0019 | YB0016 | Lula | x | | x | x | x | |
| Collection_G0019_2255 | G0019 | YB0020 | Lula | x | x | x | | | |
| Collection_G0019_2886 | G0019 | YB0003 | Lula | x | | x | | | |
| Collection_G0019_2913 | G0019 | YB0007 | Lula | x | | x | | | |
| Collection_G0020FOG_2639 | G0020 | YB0010 | Lula | x | | x | x | x | |
| Collection_G0020_1985 | G0020 | YB0021 | Lula | x | x | x | | | |
| Collection_G0020_2908 | G0020 | YB0007 | Lula | | | x | | | |
| Collection_G0020_2277 | G0020 | YB0367 | Lula | | | x | | | |
| Collection_G0020_3222 | G0020 | YB0009 | Lula | x | | x | | | |
| Collection_G0021FOG_1986 | G0021 | YB0022 | Lula | x | x | x | x | | |
| Collection_G0021_2051 | G0021 | YB0094 | Lula | x | x | x | | | |
| Collection_G0022FOG_1987 | G0022 | YB0023 | Lula | x | x | x | x | | |
| Collection_G0023FOG_1988 | G0023 | YB0024 | Lula-Wild | x | x | x | x | x | |
| Collection_G0024FOG_1989 | G0024 | YB0025 | Lula | x | x | x | x | x | |
| Collection_G0025FOG_2256 | G0025 | YB0027 | Lula | x | x | x | x | x | |
| Collection_G0026FOG_1992 | G0026 | YB0028 | Lula | x | x | x | x | x | |
| Collection_G0027FOG_1993 | G0027 | YB0029 | Lula | x | x | x | x | | |
| Collection_G0028FOG_1994 | G0028 | YB0030 | Lula-Wild | x | x | x | x | x | |
| Collection_G0029FOG_2257 | G0029 | YB0031 | Lula | x | x | x | x | x | |
| Collection_G0029_2704 | G0029 | YB0031 | Lula | x | | x | | | |
| Collection_G0030FOG_1997 | G0030 | YB0033 | Lula | x | x | x | x | | |
| Collection_G0031FOG_1998 | G0031 | YB0034 | Lula | x | x | x | x | x | |
| Collection_G0031_2718 | G0031 | YB0034 | Lula | | | x | | | |
| Collection_G0031_2722 | G0031 | YB0034 | Lula | | | x | | | |
| Collection_G0031_2282 | G0031 | YB0363 | Lula | x | | x | | | |
| Collection_G0032FOG_1999 | G0032 | YB0035 | Lula | x | x | x | x | | |
| **Unique sample name** | **Unique genetic fingerprint** | **Documented identity** | **Assigned Genotype** | **Discovery Panel** | **Validation Panel** | **Screening Panel** | **Canephora Panel** | **CC-I** | **CC-X** |

| | | | | | | | | | |
|---|---|---|---|---|---|---|---|---|---|
| Collection_G0032_3192 | G0032 | YB0035 | Lula | x | | x | | | |
| Collection_G0033FOG_2000 | G0033 | YB0036 | Lula | x | x | x | x | | |
| Collection_G0034FOG_2610 | G0034 | YB0003 | Lula | x | | x | x | | |
| Collection_G0034_2613 | G0034 | YB0003 | Lula | | | x | | | |
| Collection_G0034_2621 | G0034 | YB0007 | Lula | | | x | | | |
| Collection_G0034_2622 | G0034 | YB0007 | Lula | | | x | | | |
| Collection_G0034_2258 | G0034 | YB0037 | Lula | x | x | x | | | |
| Collection_G0034_2445 | G0034 | YB0037 | Lula | | | x | | | |
| Collection_G0034_2988 | G0034 | YB0007 | Lula | | | x | | | |
| Collection_G0034_3002 | G0034 | YB0007 | Lula | | | x | | | |
| Collection_G0034_3076 | G0034 | YB0004 | Lula | | | x | | | |
| Collection_G0035FOG_2259 | G0035 | YB0038 | Lula-Wild | x | x | x | x | | |
| Collection_G0035_2742 | G0035 | YB0038 | Lula-Wild | | | x | | | |
| Collection_G0035_3196 | G0035 | YB0038 | Lula-Wild | x | | x | | | |
| Collection_G0035_2745 | G0035 | YB0038 | Lula-Wild | | | x | | | |
| Collection_G0035_2746 | G0035 | YB0038 | Lula-Wild | | | x | | | |
| Collection_G0035_2478 | G0035 | YB0038 | Lula-Wild | | | x | | | |
| Collection_G0036FOG_2072 | G0036 | YB0039 | Lula | x | x | x | x | | |
| Collection_G0036_2003 | G0036 | YB0039 | Lula | x | x | x | | | |
| Collection_G0037FOG_2260 | G0037 | YB0040 | Lula-Wild | x | x | x | x | x | |
| Collection_G0038FOG_2005 | G0038 | YB0041 | Lula-Wild | x | x | x | x | | |
| Collection_G0039FOG_1996 | G0039 | YB0032 | Lula-subgroup A | x | x | x | x | | |
| Collection_G0039_2006 | G0039 | YB0042 | Lula-subgroup A | x | x | x | | | |
| Collection_G0040FOG_2007 | G0040 | YB0043 | Lula | x | x | x | x | x | |
| Collection_G0041FOG_2274 | G0041 | YB0013 | Lula-Wild | x | | x | x | x | |
| Collection_G0041_2008 | G0041 | YB0044 | Lula-Wild | x | x | x | | | |
| Collection_G0041_3205 | G0041 | YB0044 | Lula-Wild | x | | x | | | |
| Collection_G0042FOG_2009 | G0042 | YB0045 | Lula-Wild | x | x | x | x | | |
| Collection_G0043FOG_2010 | G0043 | YB0046 | Lula-subgroup A | x | x | x | x | x | |
| Collection_G0043_3207 | G0043 | YB0046 | Lula-subgroup A | x | | x | | | |
| Collection_G0043_2456 | G0043 | YB0046 | Lula-subgroup A | | | x | | | |
| Collection_G0043_2457 | G0043 | YB0046 | Lula-subgroup A | | | x | | | |
| Collection_G0044FOG_2011 | G0044 | YB0047 | Lula-Wild | x | x | x | x | x | |
| Collection_G0044_3209 | G0044 | YB0047 | Lula-Wild | x | | x | | | |
| Collection_G0044_3210 | G0044 | YB0047 | Lula-Wild | x | | x | | | |
| Collection_G0044_2459 | G0044 | YB0047 | Lula-Wild | x | | x | | | |
| Collection_G0044_2460 | G0044 | YB0047 | Lula-Wild | | | x | | | |
| Collection_G0045FOG_2012 | G0045 | YB0048 | Lula-Wild | | | x | x | | |
| Collection_G0046FOG_2261 | G0046 | YB0049 | Lula | x | x | x | x | | |
| Collection_G0047FOG_2014 | G0047 | YB0050 | Lula | x | x | x | x | | |
| Collection_G0048FOG_2016 | G0048 | YB0052 | Wild | x | x | x | x | | |
| Collection_G0049FOG_2017 | G0049 | YB0053 | Wild | x | x | x | x | | |
| Collection_G0050FOG_2275 | G0050 | YB0054 | Wild | x | | x | x | | |
| Collection_G0051FOG_2019 | G0051 | YB0055 | Wild | x | x | x | x | x | |
| Collection_G0052FOG_2020 | G0052 | YB0056 | Wild | x | x | x | x | x | |
| Collection_G0053FOG_2021 | G0053 | YB0057 | Wild | x | x | x | x | | |
| Collection_G0054FOG_2022 | G0054 | YB0058 | Wild | x | x | x | x | x | x |
| Collection_G0055FOG_2023 | G0055 | YB0059 | Lula | x | | x | x | | |
| Collection_G0055_2250 | G0055 | YB0059 | Lula | x | | x | | | |
| Collection_G0056FOG_2025 | G0056 | YB0061 | Lula | x | x | x | x | x | |

| Unique sample name | Unique genetic fingerprint | Documented identity | Assigned Genotype | Discovery Panel | Validation Panel | Screening Panel | Canephora Panel | CC-I | CC-X |
|---|---|---|---|---|---|---|---|---|---|
| Collection_G0057FOG_2026 | G0057 | YB0062 | Lula | x | x | x | x | | |
| Collection_G0058FOG_2027 | G0058 | YB0063 | Lula-Wild | x | x | x | x | x | |
| Collection_G0059FOG_2028 | G0059 | YB0064 | Lula-Wild | x | x | x | x | | |
| Collection_G0060FOG_2029 | G0060 | YB0065 | Lula-Wild | x | x | x | x | | |
| Collection_G0061FOG_2030 | G0061 | YB0066 | Lula-Wild | x | x | x | x | x | x |
| Collection_G0062FOG_2031 | G0062 | YB0067 | Lula-Wild | x | x | x | x | x | |
| Collection_G0063FOG_2032 | G0063 | YB0068 | Lula | x | x | x | x | x | |
| Collection_G0064FOG_2033 | G0064 | YB0069 | Lula-Wild | x | x | x | x | x | |
| Collection_G0065FOG_2034 | G0065 | YB0070 | Lula-Wild | x | x | x | x | x | |
| Collection_G0066FOG_2035 | G0066 | YB0072 | Lula-Wild | x | x | x | x | | |
| Collection_G0067FOG_2036 | G0067 | YB0073 | Lula-Wild | x | x | x | x | | |
| Collection_G0068FOG_2263 | G0068 | YB0074 | Lula | x | x | x | x | | |
| Collection_G0069FOG_2264 | G0069 | YB0077 | Lula-Wild | x | x | x | x | x | |
| Collection_G0070FOG_2265 | G0070 | YB0078 | Wild | x | x | x | x | | |
| Collection_G0071FOG_2266 | G0071 | YB0079 | Lula | x | x | x | x | | |
| Collection_G0072FOG_2041 | G0072 | YB0080 | Lula | x | x | x | x | | |
| Collection_G0073FOG_2267 | G0073 | YB0081 | Lula-subgroup A | x | x | x | x | | |
| Collection_G0074FOG_2268 | G0074 | YB0082 | Lula-subgroup A | x | x | x | x | x | |
| Collection_G0075FOG_2044 | G0075 | YB0085 | Lula | x | x | x | x | | |
| Collection_G0076FOG_2475 | G0076 | YB0086 | Lula | x | | x | x | | |
| Collection_G0077FOG_2476 | G0077 | YB0087 | Lula | x | x | x | x | x | |
| Collection_G0078FOG_2048 | G0078 | YB0090 | Lula-Wild | x | x | x | x | x | |
| Collection_G0079FOG_2049 | G0079 | YB0091 | Lula-subgroup A | x | x | x | x | x | |
| Collection_G0080FOG_2050 | G0080 | YB0092 | Lula | x | x | x | x | | |
| Collection_G0081FOG_2052 | G0081 | YB0094 | Lula-subgroup A | x | x | x | x | x | x |
| Collection_G0081_3065 | G0081 | YB0004 | Lula-subgroup A | x | | x | | | |
| Collection_G0081_3263 | G0081 | YB0094 | Lula-subgroup A | x | | x | | | |
| Collection_G0081_3260 | G0081 | YB0004 | Lula-subgroup A | x | | x | | | |
| Collection_G0082FOG_2271 | G0082 | YB0095 | Wild | x | x | x | x | x | |
| Collection_G0083FOG_2272 | G0083 | YB0096 | Lula-Wild | x | | x | x | x | |
| Collection_G0084FOG_2055 | G0084 | YB0097 | Lula | x | x | x | x | | |
| Collection_G0085FOG_2056 | G0085 | YB0098 | Lula-subgroup A | x | x | x | x | | |
| Collection_G0086FOG_2057 | G0086 | YB0357 | Lula-subgroup A | x | x | x | x | x | |
| Collection_G0087FOG_2273 | G0087 | YB0358 | Congolese subgroup A | x | x | x | x | | |
| Collection_G0088FOG_2059 | G0088 | MUTYB0002 | Lula-subgroup A | x | x | x | x | | |
| Collection_G0089FOG_2060 | G0089 | MUTYB0004 | Lula-Wild | x | | x | x | | |
| Collection_G0090FOG_2061 | G0090 | MUTYB0007A | Lula | x | x | x | x | | |
| Collection_G0091FOG_2062 | G0091 | MUTYB0007B | Lula | x | x | x | x | x | |
| Collection_G0092FOG_2063 | G0092 | MUTYB0081 | Lula-subgroup A | x | | x | x | x | |
| Collection_G0093FOG_2614 | G0093 | YB0004 | Lula | x | | x | x | | |
| Collection_G0093_2675 | G0093 | YB0003 | Lula | | | x | | | |
| Collection_G0093_2463 | G0093 | YB0001 | Lula | x | x | x | | | |
| Collection_G0093_2872 | G0093 | YB0001 | Lula | | | x | | | |
| Collection_G0093_3059 | G0093 | YB0004 | Lula | | | x | | | |
| Collection_G0093_3139 | G0093 | YB0003 | Lula | | | x | | | |
| Collection_G0093_3140 | G0093 | YB0003 | Lula | | | x | | | |
| Collection_G0093_3141 | G0093 | YB0003 | Lula | | | x | | | |
| Collection_G0094FOG_2065 | G0094 | YB0002 | Lula | x | x | x | x | x | |
| Collection_G0094_2806 | G0094 | YB0002 | Lula | | | x | | | |

| Unique sample name | Unique genetic fingerprint | Documented identity | Assigned Genotype | Discovery Panel | Validation Panel | Screening Panel | Canephora Panel | CC-I | CC-X |
|---|---|---|---|---|---|---|---|---|---|
| Collection_G0094_3271 | G0094 | YB0002 | Lula | x | | x | | | |
| Collection_G0094_2936 | G0094 | YB0002 | Lula | | | x | | | |
| Collection_G0094_2938 | G0094 | YB0002 | Lula | | | x | | | |
| Collection_G0094_2941 | G0094 | YB0002 | Lula | | | x | | | |
| Collection_G0094_2942 | G0094 | YB0002 | Lula | | | x | | | |
| Collection_G0094_2949 | G0094 | YB0002 | Lula | | | x | | | |
| Collection_G0094_2950 | G0094 | YB0002 | Lula | | | x | | | |
| Collection_G0094_2965 | G0094 | YB0002 | Lula | | | x | | | |
| Collection_G0094_2966 | G0094 | YB0002 | Lula | | | x | | | |
| Collection_G0094_2967 | G0094 | YB0002 | Lula | | | x | | | |
| Collection_G0094_2968 | G0094 | YB0002 | Lula | | | x | | | |
| Collection_G0094_2970 | G0094 | YB0002 | Lula | x | | x | | | |
| Collection_G0094_3119 | G0094 | YB0002 | Lula | x | | x | | | |
| Collection_G0095FOG_2611 | G0095 | YB0003 | Lula | x | | x | x | | |
| Collection_G0095_2620 | G0095 | YB0008 | Lula | | | x | | | |
| Collection_G0095_2623 | G0095 | YB0007 | Lula | | | x | | | |
| Collection_G0095_2739 | G0095 | YB0037 | Lula | | | x | | | |
| Collection_G0095_2894 | G0095 | YB0003 | Lula | | | x | | | |
| Collection_G0095_2898 | G0095 | YB0003 | Lula | | | x | | | |
| Collection_G0095_2914 | G0095 | YB0007 | Lula | | | x | | | |
| Collection_G0095_3009 | G0095 | YB0007 | Lula | | | x | | | |
| Collection_G0095_2343 | G0095 | YB0004 | Lula | | | x | | | |
| Collection_G0095_3070 | G0095 | YB0004 | Lula | | | x | | | |
| Collection_G0095_3071 | G0095 | YB0004 | Lula | | | x | | | |
| Collection_G0095_3124 | G0095 | YB0002 | Lula | | | x | | | |
| Collection_G0095_3146 | G0095 | YB0001 | Lula | | | x | | | |
| Collection_G0096FOG_2068 | G0096 | YB0006 | Lula | x | x | x | x | | |
| Collection_G0096_3095 | G0096 | YB0004 | Lula | | | x | | | |
| Collection_G0097FOG_2069 | G0097 | YB0006 | Lula-Wild | x | x | x | x | x | |
| Collection_G0097_2461 | G0097 | YB0047 | Lula-Wild | | | x | | | |
| Collection_G0097_3056 | G0097 | YB0005 | Lula-Wild | | | x | | | |
| Collection_G0098FOG_2910 | G0098 | YB0007 | Lula | x | | x | x | | |
| Collection_G0099FOG_2479 | G0099 | YB0059 | Lula | x | | x | x | x | |
| Collection_G0099_3217 | G0099 | YB0060 | Lula | x | | x | | | |
| Collection_G0100FOG_2076 | G0100 | YB0081 | Lula-subgroup A | x | x | x | x | x | |
| Collection_G0101FOG_2077 | G0101 | YB0082 | Lula-Wild | x | x | x | x | | |
| Collection_G0102FOG_3215 | G0102 | YB0357 | Lula-subgroup A | x | x | x | x | x | |
| Collection_G0103FOG_2604 | G0103 | YB0001 | Lula | x | x | x | x | | |
| Collection_G0103_2605 | G0103 | YB0001 | Lula | | | x | | | |
| Collection_G0103_2609 | G0103 | YB0002 | Lula | | | x | | | |
| Collection_G0103_3245 | G0103 | YB0001 | Lula | x | | x | | | |
| Collection_G0103_2866 | G0103 | YB0001 | Lula | | | x | | | |
| Collection_G0103_2874 | G0103 | YB0001 | Lula | | | x | | | |
| Collection_G0103_3024 | G0103 | YB0005 | Lula | | | x | | | |
| Collection_G0103_3225 | G0103 | YB0003 | Lula | x | | x | | | |
| Collection_G0103_3142 | G0103 | YB0003 | Lula | | | x | | | |
| Collection_G0103_3270 | G0103 | YB0001 | Lula | x | | x | | | |
| Collection_G0103_2347 | G0103 | YB0001 | Lula | | | x | | | |
| Collection_G0103_3186 | G0103 | YB0005 | Lula | | | x | | | |

| Unique sample name | Unique genetic fingerprint | Documented identity | Assigned Genotype | Discovery Panel | Validation Panel | Screening Panel | Canephora Panel | CC-I | CC-X |
|---|---|---|---|---|---|---|---|---|---|
| Collection_G0103_3255 | G0103 | YB0001 | Lula | x | | x | | | |
| Collection_G0104FOG_2606 | G0104 | YB0002 | Lula | x | | x | x | | |
| Collection_G0104_2665 | G0104 | YB0014 | Lula | | | x | | | |
| Collection_G0104_2085 | G0104 | YB0007 | Lula | x | x | x | | | |
| Collection_G0104_2977 | G0104 | YB0007 | Lula | | | x | | | |
| Collection_G0104_2981 | G0104 | YB0007 | Lula | | | x | | | |
| Collection_G0104_3047 | G0104 | YB0005 | Lula | | | x | | | |
| Collection_G0104_3131 | G0104 | YB0007 | Lula | | | x | | | |
| Collection_G0105FOG_2676 | G0105 | YB0004 | Lula | x | | x | x | | |
| Collection_G0105_3230 | G0105 | YB0004 | Lula | x | | x | | | |
| Collection_G0105_2466 | G0105 | YB0001 | Lula | | | x | | | |
| Collection_G0105_3062 | G0105 | YB0004 | Lula | | | x | | | |
| Collection_G0105_3072 | G0105 | YB0004 | Lula | x | | x | | | |
| Collection_G0105_3266 | G0105 | YB0006 | Lula | | | x | | | |
| Collection_G0105_3236 | G0105 | YB0004 | Lula | | | x | | | |
| Collection_G0105_3259 | G0105 | YB0004 | Lula | x | | x | | | |
| Collection_G0105_3250 | G0105 | YB0006 | Lula | x | x | x | | | |
| Collection_G0106FOG_2617 | G0106 | YB0004 | Lula | x | | x | x | x | |
| Collection_G0106_2628 | G0106 | YB0006 | Lula | | | x | | | |
| Collection_G0106_2677 | G0106 | YB0004 | Lula | | | x | | | |
| Collection_G0106_2678 | G0106 | YB0004 | Lula | | | x | | | |
| Collection_G0106_2897 | G0106 | YB0003 | Lula | | | x | | | |
| Collection_G0106_2918 | G0106 | YB0006 | Lula | | | x | | | |
| Collection_G0106_3014 | G0106 | YB0005 | Lula | | | x | | | |
| Collection_G0106_3018 | G0106 | YB0005 | Lula | | | x | | | |
| Collection_G0106_3035 | G0106 | YB0005 | Lula | | | x | | | |
| Collection_G0106_3044 | G0106 | YB0005 | Lula | | | x | | | |
| Collection_G0106_2083 | G0106 | YB0005 | Lula | x | x | x | | | |
| Collection_G0106_3267 | G0106 | YB0004 | Lula | x | | x | | | |
| Collection_G0106_3239 | G0106 | YB0006 | Lula | | | x | | | |
| Collection_G0106_3265 | G0106 | YB0005 | Lula | | | x | | | |
| Collection_G0107FOG_2923 | G0107 | YB0006 | Lula | x | x | x | x | | |
| Collection_G0107_2928 | G0107 | YB0006 | Lula | | | x | | | |
| Collection_G0107_3046 | G0107 | YB0005 | Lula | | | x | | | |
| Collection_G0107_3053 | G0107 | YB0005 | Lula | | | x | | | |
| Collection_G0108FOG_2278 | G0108 | YB0359 | Lula-Wild | x | | x | x | | |
| Collection_G0109FOG_2279 | G0109 | YB0360 | Lula | x | | x | x | x | |
| Collection_G0110FOG_2280 | G0110 | YB0361 | Lula | x | | x | x | x | |
| Collection_G0111FOG_2281 | G0111 | YB0362 | Lula | x | | x | x | | |
| Collection_G0112FOG_2283 | G0112 | YB0364 | Lula-Wild | x | | x | x | | |
| Collection_G0113FOG_2284 | G0113 | YB0365 | Lula | x | | x | x | | |
| Collection_G0114FOG_2285 | G0114 | YB0366 | Lula | x | | x | x | x | |
| Collection_G0115FOG_3183 | G0115 | YB0004 | Lula | x | | x | x | | |
| Collection_G0115_2348 | G0115 | YB0004 | Lula | | | x | | | |
| Collection_G0115_3185 | G0115 | YB0004 | Lula | | | x | | | |
| Collection_G0115_3249 | G0115 | YB0005 | Lula | | | x | | | |
| Collection_G0116FOG_2810 | G0116 | YB0003 | Lula | | | x | x | | |
| Collection_G0116_2975 | G0116 | YB0007 | Lula | | | x | | | |
| Collection_G0116_2994 | G0116 | YB0007 | Lula | | | x | | | |

| Unique sample name | Unique genetic fingerprint | Documented identity | Assigned Genotype | Discovery Panel | Validation Panel | Screening Panel | Canephora Panel | CC-I | CC-X |
|---|---|---|---|---|---|---|---|---|---|
| Collection_G0116_3001 | G0116 | YB0007 | Lula | | | x | | | |
| Collection_G0117FOG_3010 | G0117 | YB0007 | Lula | | | x | x | x | |
| Collection_G0117_3132 | G0117 | YB0007 | Lula | | | x | | | |
| Collection_G0118FOG_2813 | G0118 | YB0007 | Lula | | | x | x | | |
| Collection_G0118_3011 | G0118 | YB0007 | Lula | | | x | | | |
| Collection_G0118_3130 | G0118 | YB0007 | Lula | | | x | | | |
| Collection_G0119FOG_2624 | G0119 | YB0007 | Lula | x | | x | x | | |
| Collection_G0119_2625 | G0119 | YB0006 | Lula | | | x | | | |
| Collection_G0119_2627 | G0119 | YB0006 | Lula | | | x | | | |
| Collection_G0119_2640 | G0119 | YB0010 | Lula | | | x | | | |
| Collection_G0119_2642 | G0119 | YB0011 | Lula | | | x | | | |
| Collection_G0119_2782 | G0119 | YB0047 | Lula | | | x | | | |
| Collection_G0119_3211 | G0119 | YB0047 | Lula | x | | x | | | |
| Collection_G0119_2803 | G0119 | YB0002 | Lula | | | x | | | |
| Collection_G0119_3229 | G0119 | YB0006 | Lula | x | | x | | | |
| Collection_G0119_2939 | G0119 | YB0002 | Lula | | | x | | | |
| Collection_G0119_2951 | G0119 | YB0002 | Lula | | | x | | | |
| Collection_G0119_2952 | G0119 | YB0002 | Lula | | | x | | | |
| Collection_G0119_2955 | G0119 | YB0002 | Lula | | | x | | | |
| Collection_G0119_2957 | G0119 | YB0002 | Lula | | | x | | | |
| Collection_G0119_2985 | G0119 | YB0007 | Lula | | | x | | | |
| Collection_G0119_3012 | G0119 | YB0007 | Lula | | | x | | | |
| Collection_G0119_3016 | G0119 | YB0005 | Lula | | | x | | | |
| Collection_G0119_3023 | G0119 | YB0005 | Lula | | | x | | | |
| Collection_G0120FOG_2925 | G0120 | YB0006 | Lula | | | x | x | | |
| Collection_G0120_2926 | G0120 | YB0006 | Lula | | | x | | | |
| Collection_G0120_2932 | G0120 | YB0002 | Lula | | | x | | | |
| Collection_G0120_3015 | G0120 | YB0005 | Lula | | | x | | | |
| Collection_G0120_3017 | G0120 | YB0005 | Lula | | | x | | | |
| Collection_G0120_3020 | G0120 | YB0005 | Lula | | | x | | | |
| Collection_G0120_3022 | G0120 | YB0005 | Lula | | | x | | | |
| Collection_G0120_3026 | G0120 | YB0005 | Lula | | | x | | | |
| Collection_G0120_3027 | G0120 | YB0005 | Lula | | | x | | | |
| Collection_G0120_3028 | G0120 | YB0005 | Lula | | | x | | | |
| Collection_G0120_3030 | G0120 | YB0005 | Lula | | | x | | | |
| Collection_G0120_3034 | G0120 | YB0005 | Lula | | | x | | | |
| Collection_G0120_3036 | G0120 | YB0005 | Lula | | | x | | | |
| Collection_G0120_3043 | G0120 | YB0005 | Lula | | | x | | | |
| Collection_G0120_3048 | G0120 | YB0005 | Lula | | | x | | | |
| Collection_G0120_3088 | G0120 | YB0004 | Lula | | | x | | | |
| Collection_G0120_3089 | G0120 | YB0004 | Lula | | | x | | | |
| Collection_G0120_3091 | G0120 | YB0004 | Lula | | | x | | | |
| Collection_G0120_3092 | G0120 | YB0004 | Lula | | | x | | | |
| Collection_G0120_3093 | G0120 | YB0004 | Lula | | | x | | | |
| Collection_G0120_3135 | G0120 | YB0005 | Lula | | | x | | | |
| Collection_G0120_3136 | G0120 | YB0005 | Lula | | | x | | | |
| Collection_G0121FOG_2824 | G0121 | YB0005 | Lula | x | | x | x | | |
| Collection_G0121_2825 | G0121 | YB0005 | Lula | | | x | | | |
| Collection_G0121_2826 | G0121 | YB0005 | Lula | | | x | | | |

| Unique sample name | Unique genetic fingerprint | Documented identity | Assigned Genotype | Discovery Panel | Validation Panel | Screening Panel | Canephora Panel | CC-I | CC-X |
|---|---|---|---|---|---|---|---|---|---|
| Collection_G0121_3228 | G0121 | YB0005 | Lula | x | | x | | | |
| Collection_G0121_2827 | G0121 | YB0005 | Lula | | | x | | | |
| Collection_G0121_3021 | G0121 | YB0005 | Lula | x | | x | | | |
| Collection_G0122FOG_2630 | G0122 | YB0005 | Lula | | | x | x | | |
| Collection_G0122_2631 | G0122 | YB0005 | Lula | | | x | | | |
| Collection_G0122_3045 | G0122 | YB0005 | Lula | | | x | | | |
| Collection_G0123FOG_2629 | G0123 | YB0005 | Lula | | | x | x | | |
| Collection_G0123_3055 | G0123 | YB0005 | Lula | | | x | | | |
| Collection_G0123_3137 | G0123 | YB0005 | Lula | | | x | | | |
| Collection_G0124FOG_3058 | G0124 | YB0004 | Lula | | | x | x | x | |
| Collection_G0125FOG_2815 | G0125 | YB0004 | Lula | | | x | x | | |
| Collection_G0125_3060 | G0125 | YB0004 | Lula | | | x | | | |
| Collection_G0125_3061 | G0125 | YB0004 | Lula | | | x | | | |
| Collection_G0125_3068 | G0125 | YB0004 | Lula | | | x | | | |
| Collection_G0125_3074 | G0125 | YB0004 | Lula | | | x | | | |
| Collection_G0125_3084 | G0125 | YB0004 | Lula | | | x | | | |
| Collection_G0126FOG_2626 | G0126 | YB0006 | Lula-Wild | | | x | x | | |
| Collection_G0126_3067 | G0126 | YB0004 | Lula-Wild | | | x | | | |
| Collection_G0126_3187 | G0126 | YB0009 | Lula-Wild | | | x | | | |
| Collection_G0127FOG_3075 | G0127 | YB0004 | Lula | x | | x | x | | |
| Collection_G0127_3241 | G0127 | YB0002 | Lula | | | x | | | |
| Collection_G0128FOG_2892 | G0128 | YB0003 | Lula | | | x | x | | |
| Collection_G0128_2958 | G0128 | YB0002 | Lula | | | x | | | |
| Collection_G0128_2960 | G0128 | YB0002 | Lula | | | x | | | |
| Collection_G0128_2964 | G0128 | YB0002 | Lula | | | x | | | |
| Collection_G0128_3077 | G0128 | YB0004 | Lula | | | x | | | |
| Collection_G0128_3113 | G0128 | YB0002 | Lula | | | x | | | |
| Collection_G0128_3120 | G0128 | YB0002 | Lula | | | x | | | |
| Collection_G0129FOG_2851 | G0129 | YB0094 | Lula-subgroup A | | | x | x | | |
| Collection_G0129_3085 | G0129 | YB0004 | Lula-subgroup A | | | x | | | |
| Collection_G0130FOG_3096 | G0130 | YB0002 | Lula-Wild | | | x | x | | |
| Collection_G0131FOG_2919 | G0131 | YB0006 | Lula | | | x | x | | |
| Collection_G0131_2933 | G0131 | YB0002 | Lula | | | x | | | |
| Collection_G0131_2945 | G0131 | YB0002 | Lula | | | x | | | |
| Collection_G0131_2954 | G0131 | YB0002 | Lula | | | x | | | |
| Collection_G0131_2959 | G0131 | YB0002 | Lula | | | x | | | |
| Collection_G0131_3097 | G0131 | YB0002 | Lula | | | x | | | |
| Collection_G0131_3099 | G0131 | YB0002 | Lula | | | x | | | |
| Collection_G0131_3106 | G0131 | YB0002 | Lula | | | x | | | |
| Collection_G0131_3115 | G0131 | YB0002 | Lula | | | x | | | |
| Collection_G0131_3121 | G0131 | YB0002 | Lula | | | x | | | |
| Collection_G0131_3122 | G0131 | YB0002 | Lula | | | x | | | |
| Collection_G0132FOG_2805 | G0132 | YB0002 | Lula | | | x | x | | |
| Collection_G0132_2937 | G0132 | YB0002 | Lula | | | x | | | |
| Collection_G0132_2948 | G0132 | YB0002 | Lula | | | x | | | |
| Collection_G0132_2969 | G0132 | YB0002 | Lula | | | x | | | |
| Collection_G0132_3098 | G0132 | YB0002 | Lula | | | x | | | |
| Collection_G0132_3102 | G0132 | YB0002 | Lula | | | x | | | |
| Collection_G0132_3109 | G0132 | YB0002 | Lula | | | x | | | |

| Unique sample name | Unique genetic fingerprint | Documented identity | Assigned Genotype | Discovery Panel | Validation Panel | Screening Panel | Canephora Panel | CC-I | CC-X |
|---|---|---|---|---|---|---|---|---|---|
| Collection_G0133FOG_2963 | G0133 | YB0002 | Lula | | | x | x | | |
| Collection_G0133_3104 | G0133 | YB0002 | Lula | | | x | | | |
| Collection_G0134FOG_2807 | G0134 | YB0002 | Lula | x | | x | x | x | |
| Collection_G0134_2953 | G0134 | YB0002 | Lula | | | x | | | |
| Collection_G0134_2956 | G0134 | YB0002 | Lula | | | x | | | |
| Collection_G0134_2962 | G0134 | YB0002 | Lula | | | x | | | |
| Collection_G0134_3233 | G0134 | YB0002 | Lula | | | x | | | |
| Collection_G0134_3108 | G0134 | YB0002 | Lula | | | x | | | |
| Collection_G0134_3118 | G0134 | YB0002 | Lula | | | x | | | |
| Collection_G0135FOG_2830 | G0135 | YB0007 | Lula | x | | x | x | x | |
| Collection_G0135_2961 | G0135 | YB0002 | Lula | | | x | | | |
| Collection_G0135_2971 | G0135 | YB0002 | Lula | | | x | | | |
| Collection_G0135_3110 | G0135 | YB0002 | Lula | | | x | | | |
| Collection_G0135_3181 | G0135 | YB0002 | Lula | | | x | | | |
| Collection_G0135_3182 | G0135 | YB0002 | Lula | | | x | | | |
| Collection_G0135_3252 | G0135 | YB0007 | Lula | | | x | | | |
| Collection_G0136FOG_3134 | G0136 | YB0005 | Lula | | | x | x | | |
| Collection_G0137FOG_3138 | G0137 | YB0005 | Lula | | | x | x | | |
| Collection_G0138FOG_3231 | G0138 | YB0007 | Lula | x | | x | x | | |
| Collection_G0138_3244 | G0138 | YB0007 | Lula | x | | x | | | |
| Collection_G0138_3147 | G0138 | YB0001 | Lula | | | x | | | |
| Collection_G0139FOG_3150 | G0139 | YB0081 | Lula-subgroup A | | | x | x | | |
| Collection_G0140FOG_3151 | G0140 | YB0081 | Lula-subgroup A | | | x | x | x | |
| Collection_G0141FOG_3152 | G0141 | YB0081 | Lula | | | x | x | | |
| Collection_G0142FOG_3153 | G0142 | YB0081 | Lula-subgroup A | | | x | x | x | |
| Collection_G0143FOG_3154 | G0143 | YB0081 | Lula | | | x | x | | |
| Collection_G0144FOG_3162 | G0144 | YB0081 | Lula-subgroup A | | | x | x | | |
| Collection_G0145FOG_3163 | G0145 | YB0081 | Lula-subgroup A | | | x | x | | |
| Collection_G0146FOG_3157 | G0146 | YB0081 | Lula-subgroup A | | | x | x | | |
| Collection_G0147FOG_3158 | G0147 | YB0081 | Lula-subgroup A | | | x | x | x | |
| Collection_G0148FOG_3159 | G0148 | YB0081 | Lula-subgroup A | | | x | x | | |
| Collection_G0149FOG_3160 | G0149 | YB0081 | Lula-subgroup A | | | x | x | | |
| Collection_G0150FOG_3161 | G0150 | YB0081 | Lula-subgroup A | | | x | x | | x |
| Collection_G0151FOG_3155 | G0151 | YB0366 | Lula-Wild | | | x | x | x | |
| Collection_G0152FOG_3156 | G0152 | YB0366 | Lula | | | x | x | x | |
| Collection_G0153FOG_3164 | G0153 | YB0359 | Lula-Wild | | | x | x | | x |
| Collection_G0154FOG_3166 | G0154 | YB0360 | Lula | | | x | x | | |
| Collection_G0155FOG_3167 | G0155 | YB0361 | Lula-Wild | | | x | x | | |
| Collection_G0156FOG_3168 | G0156 | YB0361 | Lula | | | x | x | | |
| Collection_G0157FOG_3170 | G0157 | YB0362 | Lula | | | x | x | x | |
| Collection_G0158FOG_3178 | G0158 | YB0365 | Lula | | | x | x | x | |
| Collection_G0159FOG_3172 | G0159 | YB0363 | Lula | | | x | x | | |
| Collection_G0160FOG_3173 | G0160 | YB0363 | Lula | | | x | x | | |
| Collection_G0161FOG_3174 | G0161 | YB0364 | Lula | | | x | x | | |
| Collection_G0162FOG_3175 | G0162 | YB0364 | Lula | | | x | x | x | |
| Collection_G0163FOG_3176 | G0163 | YB0364 | Lula | | | x | x | | |
| Collection_G0164FOG_3177 | G0164 | YB0365 | Lula | | | x | x | | |
| Collection_G0165FOG_3191 | G0165 | YB0034 | Lula | | | x | x | | |
| Collection_G0165_2720 | G0165 | YB0034 | Lula | | | x | | | |

| Unique sample name | Unique genetic fingerprint | Documented identity | Assigned Genotype | Discovery Panel | Validation Panel | Screening Panel | Canephora Panel | CC-I | CC-X |
|---|---|---|---|---|---|---|---|---|---|
| Collection_G0165_2721 | G0165 | YB0034 | Lula | | | x | | | |
| Collection_G0165_2723 | G0165 | YB0034 | Lula | | | x | | | |
| Collection_G0165_3171 | G0165 | YB0363 | Lula | | | x | | | |
| Collection_G0166FOG_2657 | G0166 | YB0001 | Lula-subgroup A | | | x | x | x | |
| Collection_G0167FOG_2666 | G0167 | YB0002 | Lula | | | x | x | x | |
| Collection_G0168FOG_2673 | G0168 | YB0003 | Lula | | | x | x | | |
| Collection_G0169FOG_2615 | G0169 | YB0004 | Lula | | | x | x | x | |
| Collection_G0169_2674 | G0169 | YB0003 | Lula | | | x | | | |
| Collection_G0170FOG_2632 | G0170 | YB0005 | Lula | | | x | x | | |
| Collection_G0171FOG_2634 | G0171 | YB0009 | Lula | | | x | x | | |
| Collection_G0172FOG_2655 | G0172 | YB0016 | Lula | | | x | x | x | |
| Collection_G0173FOG_2652 | G0173 | YB0015 | Lula | | | x | x | | |
| Collection_G0174FOG_2656 | G0174 | YB0001 | Lula | | | x | x | | |
| Collection_G0175FOG_2645 | G0175 | YB0013 | Lula | x | | x | x | x | x |
| Collection_G0175_2646 | G0175 | YB0013 | Lula | | | x | | | |
| Collection_G0175_2647 | G0175 | YB0013 | Lula | | | x | | | |
| Collection_G0175_2659 | G0175 | YB0017 | Lula | | | x | | | |
| Collection_G0175_3206 | G0175 | YB0013 | Lula | x | | x | | | |
| Collection_G0175_2454 | G0175 | YB0013 | Lula | | | x | | | |
| Collection_G0175_2455 | G0175 | YB0013 | Lula | | | x | | | |
| Collection_G0176FOG_3199 | G0176 | YB0040 | Lula-Wild | | | x | x | x | |
| Collection_G0177FOG_3201 | G0177 | YB0041 | Lula-Wild | | | x | x | x | |
| Collection_G0178FOG_2698 | G0178 | YB0031 | Lula | | | x | x | | |
| Collection_G0178_2700 | G0178 | YB0031 | Lula | | | x | | | |
| Collection_G0179FOG_2699 | G0179 | YB0031 | Lula-Wild | | | x | x | | |
| Collection_G0180FOG_2701 | G0180 | YB0031 | Lula-Wild | | | x | x | x | x |
| Collection_G0181FOG_2702 | G0181 | YB0031 | Lula-Wild | | | x | x | | |
| Collection_G0182FOG_2703 | G0182 | YB0031 | Lula | | | x | x | | |
| Collection_G0183FOG_2705 | G0183 | YB0032 | Lula | | | x | x | | |
| Collection_G0183_2709 | G0183 | YB0032 | Lula | | | x | | | |
| Collection_G0183_2710 | G0183 | YB0032 | Lula | | | x | | | |
| Collection_G0184FOG_2706 | G0184 | YB0032 | Lula-Wild | | | x | x | | |
| Collection_G0184_2707 | G0184 | YB0032 | Lula-Wild | | | x | | | |
| Collection_G0184_2708 | G0184 | YB0032 | Lula-Wild | | | x | | | |
| Collection_G0185FOG_2711 | G0185 | YB0032 | Lula-Wild | | | x | x | | |
| Collection_G0186FOG_2712 | G0186 | YB0033 | Lula | | | x | x | | |
| Collection_G0186_2713 | G0186 | YB0033 | Lula | | | x | | | |
| Collection_G0186_2715 | G0186 | YB0033 | Lula | | | x | | | |
| Collection_G0186_2716 | G0186 | YB0033 | Lula | | | x | | | |
| Collection_G0186_2349 | G0186 | YB0035 | Lula | | | x | | | |
| Collection_G0187FOG_2717 | G0187 | YB0033 | Lula-Wild | | | x | x | x | |
| Collection_G0188FOG_2724 | G0188 | YB0035 | Lula | | | x | x | | |
| Collection_G0188_2728 | G0188 | YB0035 | Lula | | | x | | | |
| Collection_G0189FOG_2727 | G0189 | YB0035 | Lula | | | x | x | x | |
| Collection_G0190FOG_2736 | G0190 | YB0037 | Lula-Wild | | | x | x | x | |
| Collection_G0191FOG_2741 | G0191 | YB0037 | Lula | | | x | x | x | |
| Collection_G0192FOG_2447 | G0192 | YB0038 | Lula-Wild | | | x | x | x | |
| Collection_G0192_2747 | G0192 | YB0038 | Lula-Wild | | | x | | | |
| Collection_G0193FOG_2784 | G0193 | YB0004 | Lula | | | x | x | | |

| Unique sample name | Unique genetic fingerprint | Documented identity | Assigned Genotype | Discovery Panel | Validation Panel | Screening Panel | Canephora Panel | CC-I | CC-X |
|---|---|---|---|---|---|---|---|---|---|
| Collection_G0193_2785 | G0193 | YB0004 | Lula | | | x | | | |
| Collection_G0193_2786 | G0193 | YB0004 | Lula | | | x | | | |
| Collection_G0193_2789 | G0193 | YB0004 | Lula | | | x | | | |
| Collection_G0193_2790 | G0193 | YB0004 | Lula | | | x | | | |
| Collection_G0194FOG_2730 | G0194 | YB0036 | Lula | | | x | x | | |
| Collection_G0194_3193 | G0194 | YB0036 | Lula | | | x | | | |
| Collection_G0194_2732 | G0194 | YB0036 | Lula | | | x | | | |
| Collection_G0194_2443 | G0194 | YB0036 | Lula | | | x | | | |
| Collection_G0195FOG_3194 | G0195 | YB0036 | Lula | | | x | x | x | |
| Collection_G0196FOG_3198 | G0196 | YB0039 | Lula | | | x | x | | |
| Collection_G0197FOG_3232 | G0197 | YB0013 | Lula | x | | x | x | | |
| Collection_G0197_2809 | G0197 | YB0003 | Lula | | | x | | | |
| Collection_G0198FOG_2800 | G0198 | YB0001 | Lula | x | | x | x | | |
| Collection_G0198_2863 | G0198 | YB0001 | Lula | | | x | | | |
| Collection_G0198_2465 | G0198 | YB0001 | Lula | x | | x | | | |
| Collection_G0198_2867 | G0198 | YB0001 | Lula | | | x | | | |
| Collection_G0198_2868 | G0198 | YB0001 | Lula | | | x | | | |
| Collection_G0199FOG_2801 | G0199 | YB0001 | Lula | | | x | x | | |
| Collection_G0200FOG_2802 | G0200 | YB0001 | Lula | | | x | x | x | |
| Collection_G0201FOG_2811 | G0201 | YB0003 | Lula | | | x | x | x | |
| Collection_G0202FOG_2812 | G0202 | YB0003 | Lula-Wild | | | x | x | | |
| Collection_G0203FOG_2814 | G0203 | YB0007 | Lula | | | x | x | | |
| Collection_G0203_2891 | G0203 | YB0003 | Lula | | | x | | | |
| Collection_G0204FOG_2816 | G0204 | YB0004 | Lula | | | x | x | x | |
| Collection_G0205FOG_2817 | G0205 | YB0004 | Lula | | | x | x | x | |
| Collection_G0206FOG_2818 | G0206 | YB0004 | Lula | | | x | x | | |
| Collection_G0206_2852 | G0206 | YB0094 | Lula | | | x | | | |
| Collection_G0207FOG_2820 | G0207 | YB0006 | Lula-Wild | | | x | x | x | |
| Collection_G0207_2821 | G0207 | YB0006 | Lula-Wild | | | x | | | |
| Collection_G0207_2917 | G0207 | YB0006 | Lula-Wild | | | x | | | |
| Collection_G0208FOG_2828 | G0208 | YB0005 | Lula-Wild | | | x | x | | |
| Collection_G0209FOG_2831 | G0209 | YB0358 | Congolese subgroup A | | | x | x | x | |
| Collection_G0210FOG_2841 | G0210 | YB0082 | Congolese subgroup A | | | x | x | | |
| Collection_G0211FOG_2840 | G0211 | YB0358 | Lula-subgroup A | | | x | x | x | |
| Collection_G0212FOG_2838 | G0212 | YB0357 | Lula-subgroup A | | | x | x | x | |
| Collection_G0213FOG_2833 | G0213 | YB0358 | Lula-subgroup A | | | x | x | x | |
| Collection_G0214FOG_2832 | G0214 | YB0358 | Lula-subgroup A | | | x | x | x | |
| Collection_G0215FOG_2839 | G0215 | YB0357 | Lula-subgroup A | | | x | x | | |
| Collection_G0216FOG_2844 | G0216 | YB0082 | Lula-subgroup A | | | x | x | | |
| Collection_G0217FOG_2843 | G0217 | YB0082 | Lula-subgroup A | | | x | x | x | |
| Collection_G0218FOG_2836 | G0218 | YB0357 | Lula-subgroup A | | | x | x | | |
| Collection_G0219FOG_2837 | G0219 | YB0357 | Lula-subgroup A | | | x | x | | |
| Collection_G0220FOG_2846 | G0220 | YB0081 | Lula-subgroup A | | | x | x | | |
| Collection_G0221FOG_2847 | G0221 | YB0081 | Lula-subgroup A | | | x | x | x | |
| Collection_G0222FOG_2834 | G0222 | YB0357 | Lula-subgroup A | | | x | x | | |
| Collection_G0223FOG_2845 | G0223 | YB0081 | Lula-subgroup A | | | x | x | x | |
| Collection_G0224FOG_3218 | G0224 | YB0001 | Lula | | | x | x | | |
| Collection_G0225FOG_2930 | G0225 | YB0006 | Lula | | | x | x | x | |
| Collection_G0226FOG_2931 | G0226 | YB0002 | Lula | | | x | x | | |

| Unique sample name | Unique genetic fingerprint | Documented identity | Assigned Genotype | Discovery Panel | Validation Panel | Screening Panel | Canephora Panel | CC-I | CC-X |
|---|---|---|---|---|---|---|---|---|---|
| Collection_G0227FOG_2921 | G0227 | YB0006 | Lula | | | x | x | x | |
| Collection_G0228FOG_2862 | G0228 | YB0001 | Lula | | | x | x | | |
| Collection_G0229FOG_2920 | G0229 | YB0006 | Lula-Wild | | | x | x | | |
| Collection_G0229_2922 | G0229 | YB0006 | Lula-Wild | | | x | | | |
| Collection_G0230FOG_2859 | G0230 | YB0061 | Lula | x | | x | x | x | |
| Collection_G0231FOG_2856 | G0231 | YB0059 | Lula | | | x | x | x | |
| Collection_G0232FOG_2480 | G0232 | YB0060 | Lula-Wild | x | | x | x | | |
| Collection_G0233FOG_3179 | G0233 | YB0001 | Lula | x | | x | x | x | |
| Collection_G0234FOG_3238 | G0234 | YB0004 | Lula | x | | x | x | | |
| Collection_G0235FOG_2792 | G0235 | YB0049 | Lula | x | | x | x | | |
| Collection_G0236FOG_2793 | G0236 | YB0049 | Lula-Wild | | | x | x | | |
| Collection_G0237FOG_2794 | G0237 | YB0049 | Lula | | | x | x | | |
| Collection_G0238FOG_2798 | G0238 | YB0051 | Lula | | | x | x | | |
| Collection_G0239FOG_3212 | G0239 | YB0050 | Lula | | | x | x | | |
| Collection_G0240FOG_3214 | G0240 | YB0050 | Lula | | | x | x | | |
| Collection_G0241FOG_2853 | G0241 | YB0094 | Lula-Wild | | | x | x | | |
| Collection_G0241_2854 | G0241 | YB0094 | Lula-Wild | | | x | | | |
| Collection_G0242FOG_2944 | G0242 | YB0002 | Lula | | | x | x | | |
| Collection_G0243FOG_2974 | G0243 | YB0007 | Lula | | | x | x | | |
| Collection_G0244FOG_2899 | G0244 | YB0007 | Lula | | | x | x | | |
| Collection_G0244_2983 | G0244 | YB0007 | Lula | | | x | | | |
| Collection_G0245FOG_2940 | G0245 | YB0002 | Lula | x | | x | x | | |
| Collection_G0245_3221 | G0245 | YB0009 | Lula | | | x | | | |
| Collection_G0246FOG_3220 | G0246 | YB0009 | Lula | x | | x | x | | |
| Collection_G0247FOG_2467 | G0247 | YB0001 | Lula | x | | x | x | | |
| Collection_G0247_3248 | G0247 | YB0005 | Lula | | | x | | | |
| Collection_G0247_3247 | G0247 | YB0005 | Lula | | | x | | | |
| Collection_G0248FOG_3246 | G0248 | YB0003 | Lula | x | | x | x | x | |
| Collection_G0249FOG_2787 | G0249 | YB0004 | Lula | | | x | x | x | x |
| Collection_G0249_2791 | G0249 | YB0004 | Lula | | | x | | | |
| Collection_G0250FOG_1470 | G0250 | YB0043 | Lula | x | | x | x | | |
| Collection_G0251FOG_1472 | G0251 | YB0082 | Lula-subgroup A | x | | x | x | | |
| Collection_G0252FOG_1473 | G0252 | YB0357 | Lula-subgroup A | x | | x | x | | |
| Collection_G0253FOG_1474 | G0253 | YB0358 | Lula-subgroup A | x | | x | x | x | |
| Collection_G0254FOG_1475 | G0254 | YB0081 | Congolese subgroup A | x | | x | x | x | x |
| Collection_G0255FOG_1494 | G0255 | YB0094 | Lula | x | | x | x | x | x |
| Collection_G0256FOG_1498 | G0256 | YB0051 | Lula | x | | x | x | | |
| Collection_G0257FOG_2446 | G0257 | YB0037 | Lula | | | x | x | | |
| Collection_G0258FOG_2462 | G0258 | YB0004 | Lula | | | x | x | | |
| Collection_G0259FOG_2350 | G0259 | YB0082 | Lula-subgroup A | | | x | x | | |
| Collection_G0260FOG_2345 | G0260 | YB0360 | Lula | | | x | x | | |
| Collection_G0261FOG_2346 | G0261 | YB0362 | Lula | | | x | x | x | |
| Collection_G0262FOG_2477 | G0262 | YB0037 | Lula | | | x | x | x | |
| Collection_G0263FOG_2444 | G0263 | YB0036 | Lula | | | x | x | | |
